# Digital light processing of human-scale tissue-engineered heart valves

**DOI:** 10.64898/2026.11.22.768423

**Authors:** Arman Jafari, Alireza Nemati, Arman Sarshoghi, Naimeh Rafatian, Sandrine Guenette, Yeojin Park, Vahid Karamzadeh, Ali Hedayatnia, Charles Leduc, Moishe Liberman, Ismail Bouhout, Lyes Kadem, Gregor Andelfinger, Houman Savoji

**Author notes:** Corresponding author: Houman Savoji.

## Abstract

Tissue engineered heart valves (TEHVs) could overcome the complications of current heart valve replacement, such as immunologic events, degenerative fibrocalcific changes, and patient-prosthesis mismatch. Digital light processing (DLP) 3D printing enables creation of complex geometries; however, suitable (bio)inks remain limited. Here, a composite ink comprising poly(ethylene glycol) diacrylate (PEGDA), gelatin methacryloyl (GelMA), and methacrylated quince seed mucilage (QSGM), termed GPQuin, was developed for DLP printing of human-scale TEHVs. GPQuin enabled fabrication of a trileaflet TEHV with separated leaflets and anatomically inspired sinuses. Micro-computed tomography revealed mean positive and negative surface deviations of +214 and −248 µm, respectively. The printed hydrogels exhibited a tensile modulus of 117.2 ± 15.5 kPa, an ultimate tensile strength of 113.2 ± 17.4 kPa, and an elongation at break of 75.3 ± 2.3%. Human induced pluripotent stem cell-derived valve cells attached to the hydrogel surfaces and remained predominantly viable over 14 days. After 12 weeks of subcutaneous implantation, the constructs showed a time-dependent inflammatory response. In a pulse duplicator under pulmonary loading, printed valves exhibited good cyclic leaflet opening and closing. Together, these findings establish GPQuin as a printable platform for human-scale valve scaffolds and support further investigation in heart valve tissue engineering.

## 1. Introduction

Pediatric pulmonary valve replacement remains challenging because of the anatomical complexity of congenital abnormalities involving the pulmonary valve and right ventricular outflow tract (RVOT). In the United States alone, more than 130,000 pediatric patients live with pulmonary valve disease ^[1]^. Congenital abnormalities affecting the pulmonary valve and RVOT can cause outflow obstruction, pulmonary regurgitation, or both, potentially requiring surgical intervention and valve replacement ^[1]^. Conventional replacement options have important limitations in pediatric patients, including thromboembolic complications associated with mechanical valves and structural degeneration of bioprosthetic valves ^[2]^. Selecting an appropriately sized prosthesis presents an additional challenge, as excessive oversizing has been associated with earlier structural deterioration following pulmonary valve replacement ^[1]^. These limitations underscore the need for valve substitutes that accommodate pediatric RVOT anatomy while maintaining adequate mechanical and hemodynamic performance under pulmonary circulatory loading. Tissue-engineering approaches offer a potential strategy to address these requirements through scaffolds with controlled geometry and material properties designed to maintain mechanical function while potentially supporting tissue integration and remodeling ^[3]^.

Since the introduction of tissue engineering principles, this approach has been extensively explored to develop innovative solutions with fewer complications for tissue repair and replacement ^[4]^. In the case of tissue-engineered heart valves (TEHVs), the ultimate goal is to fabricate valves that are readily available, function effectively, resemble the native valve structure, possess the capacity for remodeling and growth, and do not induce immunogenicity, calcification, or thrombogenicity ^[5]^. Untill now different groups have developed TEHVs to optimize one or more of these requirements. For instance, Li et al. recently developed a a novel crosslinking strategy to regulate the immune microenvironment for their TEHV ^[6]^. Their TEHV demonstrated anti-thrombogenic and anti-calcification properties, as well as the potential for recellularization. Despite these promising outcomes, the study relied on decellularized heart valves (DHVs), which may still pose risks of immunogenicity or disease transmission. Additionally, the availability of such valves remains limited due to the dependence on donor-derived tissues. Polymer-based TEHVs present a promising alternative to DHVs, offering greater design flexibility and enabling the development of more accessible and customizable valve replacements.

To date, various techniques, including molding, electrospinning, melt electrowriting, 3D printing, and self-assembly, have been employed to fabricate polymer-based tissue-engineered scaffolds (TESs) and TEHVs ^[7]^. Among these, 3D printing has emerged as a prominent choice for fabricating TESs, as it combines high flexibility with design freedom and precise control ^[8]^. As a result, 3D printing enables the fabrication of complex structures with micron-scale features while maintaining relative ease of production. Furthermore, 3D printing facilitates the development of patient-specific structures tailored to individual anatomical and functional requirements ^[9]^. Leveraging this technique, different groups have fabricated TEHVs^[10]^. For example, we previously developed an ink suitable for 3D printing of TEHVs using extrusion-based 3D printing ^[11]^. We have successfully fabricated TEHVs of various sizes and geometries, demonstrating functional performance under physiological conditions ^[11]^. Additionally, the materials exhibited bio- and hemocompatibility with the potential for in vivo remodeling over time. However, a limitation of extrusion-based 3D printing is its relatively low resolution, as the material must be extruded through a nozzle typically measuring several hundred micrometers in diameter.

Digital light processing (DLP) constructs three-dimensional objects through spatially controlled photopolymerization of successive layers. ^[12]^. The application of DLP for fabrication of TEHVs requires formulations that preserve thin, separated leaflet geometries while providing sufficient compliance for opening and closing. Previous light-based fabrication studies have demonstrated artificial valve structures, including heterogeneous hydrogel constructs and grayscale-DLP valve models ^[13]^. However, these studies were primarily focused on early demonstration of manufacturing feasibility rather than developing biologically validated TEHV scaffolds. Nevertheless, these developments motivate further investigation of photocurable formulations to formulate an ink for fabrication of functional TEHV. In this work, we developed a novel multi-material bioink (GPQuin) composed of gelatin methacryloyl (GelMA), poly(ethylene glycol) diacrylate (PEGDA) of two molecular weights, and methacrylated quince seed mucilage (QSGM) for DLP 3D printing applications. The ink was specifically engineered to fabricate constructs with varying geometries, from simple to highly complex, while maintaining high resolution and accuracy. Additionally, its capability to fabricate geometrically intricate, human-scale constructs with tissue-specific features was investigated through the 3D printing of TEHVs (**Figure 1**). Building on our previous studies of extrusion-printed valves and photocrosslinkable quince seed mucilage ^[11, 14]^, we examine the formulation through geometric, physicochemical, mechanical, cellular, and host-response assessments. We further investigate leaflet motion and pressure waveforms in a pulmonary pulse-duplicator configuration. The study connects composite formulation and DLP fabrication with biological characterization and preliminary valve actuation.

**Figure 1.**
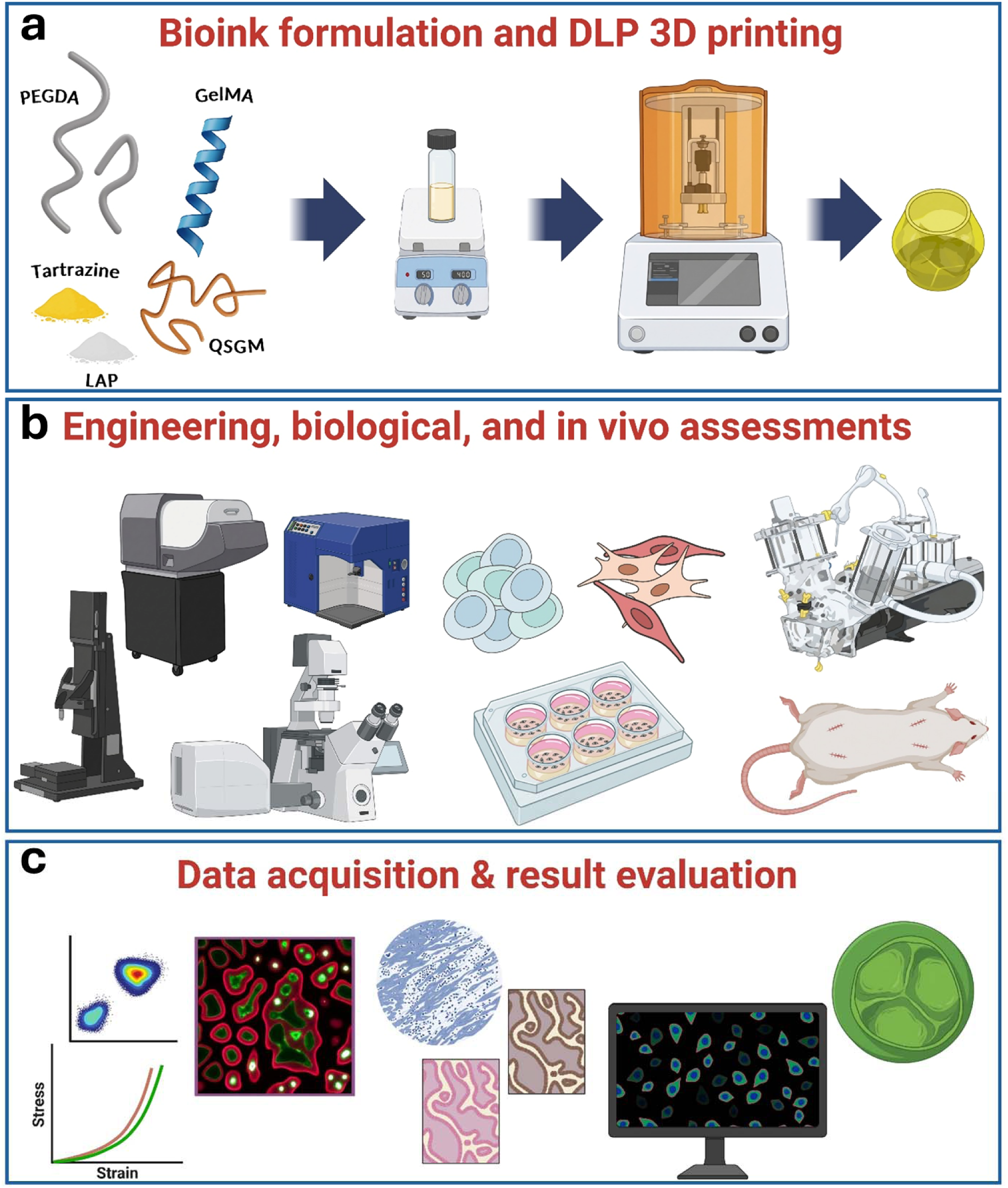
Overview of GPQuin formulation, digital light processing fabrication, and experimental evaluation of TEHVs. (a) GPQuin is formulated by dissolving naturally derived polymers (GelMA and QS-GM) and synthetic polymers (PEGDA with M_w_ of 20,000 and 700 Da) in water or cell culture medium, together with the photoinitiator (LAP) and the photoabsorber (tartrazine). This composite ink was utilized for the fabrication of various TESs, including TEHVs. (b) The physical, mechanical, and biological characteristics of GPQuin and subsequent 3D printed hydrogels were recorded and (c) the collected data was leveraged to analyze the suitability of the GPQuin hydrogels for tissue engineering applications, specifically for TEHVs.

## 2. Materials and methods

### 2.1. Synthesis of GelMA

For GelMA synthesis, 10 g of gelatin (from porcine skin, Type A, Sigma-Aldrich) was dissolved in 200 mL of phosphate buffered saline (PBS, pH = 7.4, Gibco) heated at 60 °C. Upon reaching a clear solution, 8 mL of methacrylic anhydride (MA, Sigma-Aldrich) was added dropwise to the gelatin solution. The MA/gelatin mixture was stirred for 3 h. To purify the GelMA, the solution was then dialyzed against water for 7 days using a 12-14 kDa molecular weight cut-off dialysis tube (Spectrum Labs) followed by lyophilization for 5 days. The GelMA was then stored at -20 °C until further use.

### 2.2. Extraction of QS and synthesis of QSGM

Quince seed mucilage (QS) was extracted and purified following previously described protocols by our team ^[14]^. Briefly, the seeds were immersed in ultra-pure water for 24 h under continuous stirring at 40 °C. The extracted QS was separated from the seeds via centrifugation at 4000 rpm for 10 min, and the resulting supernatant was frozen and lyophilized for 5 days to yield purified QS.

To synthesize QSGM, the QS was dissolved in ultrapure water at a concentration of 0.5 wt.%. The pH of the solution was adjusted to around 3.5 using HCl solution. To prevent undesired reactions, 4-Methoxyphenol (MEHQ, Sigma-Aldrich) was added at a concentration of 0.5 wt.% to all reactants. Glycidyl methacrylate (GM, Sigma-Aldrich) was gradually added dropwise to the solution at a GM/QS ratio of 5:1 (w/v). The reaction was allowed to proceed for 24 h while maintaining the temperature at 60 °C. Upon completion of the reaction, the solution was dialyzed and lyophilized as described above. The resulting QSGM was stored at – 20 °C until further use.

### 2.3. Formulation of the GPQuin

GPQuin was made of a mixture of natural and synthetic polymers. In a typical procedure, 0.4 g of GelMA, 0.5 g of PEGDA (M_n_ 20,000, Bezwada Biomedical), 0.3 g of PEGDA (M_n_ 700, Sigma-Aldrich), and 0.05 g of QSGM were weighed and added to 10 mL of DI water. The mixture was then heated at 37 °C and vigorously mixed to ensure complete dissolution of the polymers. To complete the inks, 0.6 mM of tartrazine (Sigma-Aldrich) and 0.06 g of lithium phenyl-2,4,6-trimethylbenzoylphosphinate (LAP, Sigma-Aldrich) were added to the mixture. The inks were kept at 37 °C until used for the printing.

### 2.4. DLP 3D printing of the constructs and printing fidelity

The 3D printing was performed using a digital light processing (DLP) 3D bioprinter (Lumen X+, Cellink, USA). The prepared inks were poured into Petri dishes coated with polydimethylsiloxane (PDMS), serving as the vat. The 3D design was transferred to the 3D bioprinter to create the printing files. The trileaflet valve geometry was obtained from the NIH 3D Print Exchange (3DPX-000452; https://3d.nih.gov/entries/3dpx-000452). This generic semilunar valve design was used as a geometric template for DLP valve fabrication. The printing parameters, including projector power level, base and body exposure times, and layer thickness, were optimized through a systematic trial-and-error analysis. Upon completion of the printing process, the printed constructs were removed from the build platform and either used for further analysis or stored in PBS as needed.

To assess the printing fidelity, two approaches were used. First, we used our own design containing features of different sizes. The design was used to find the optimum printing parameters and finding the smallest micron-sized features possible to print with the developed inks. The printing fidelity of the heart valves were evaluated by overlaying the design and printed construct. After printing, micro-CT scanning (µCT) of the valves was performed using a SkyScan1176 scanner (Bruker, Belgium). A 360° scan was acquired with a 0.7° rotation step, 2 averaging frames, a source voltage of 70 kV, a source current of 85 µA, a pixel size of 35 µm, and an exposure time of 100 ms. The scans were used to generate 3D reconstructions of the printed valves. The resulting STL files were imported into Geomagic Wrap (3D Systems) along with the original STL file used for printing. These files were overlaid, and deviations between them were quantified and visualized using heatmaps.

### 2.5. Swelling and degradation

After printing, hydrogels were washed with PBS to ensure the removal of unreacted inks. Equilibrium swelling was determined by calculating the difference between the swollen and dry weights of the hydrogels using the following equation:

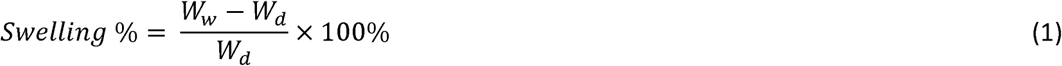

Where W_w_ represents the weight of the hydrogels after 24 h immersion in PBS at 37 °C, and W_d_ is the dry weight of the same sample.

Hydrogel degradation was assessed through hydrolytic and enzymatic degradation tests. Prior to testing, samples were dried and weighed (W_0_). For hydrolytic degradation, samples were immersed in PBS. Hydrogels were maintained at 37 °C, and at designated time points, samples were removed, washed with distilled water, and dried until a constant weight (W_t_) was achieved. The degradation medium was refreshed weekly.

For enzymatic degradation, collagenase Type II (17101-015, Gibco™, USA) was dissolved in PBS containing 5 mM CaCl₂ to achieve a final activity of 50 U/mL. Each hydrogel was incubated in collagenase containing PBS at 37 °C. Degradation was then calculated at predetermined timepoints of 3, 6, 12, 24, 48, 72, and 120 hours. At each timepoint, hydrogels were removed from the solution, rinsed in PBS, followed by immersion in 50 mM EDTA in PBS for 10 minutes at room temperature to chelate Ca²⁺ and inactivate collagenase ^[15]^. The scaffolds were then rinsed 2–3 times with deionized water, air-dried at room temperature until a stable mass was achieved (W_t_). The degradation percentage was calculated using Equation 2:

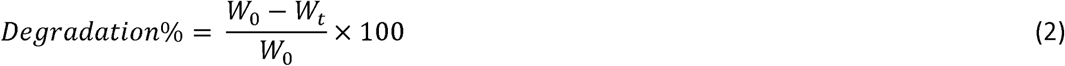

For these experiments, 3D printed hydrogels from 4% GelMA served as the control.

### 2.6. Mechanical characterization

The mechanical properties of hydrogels were characterized using a Mach-1 mechanical tester (Biomomentum, Canada). For tensile tests, rectangular samples of 30 × 6 × 1 mm^3^ were printed using optimized printing parameters (n = 5), mounted in grips, and were pulled at the rate of 10 mm.min^−1^. For the compression tests, cylindrical samples (10 mm in diameter, 5 mm in height) were similarly printed (n = 5). Samples were then compressed at the rate of 5 mm.min^−1^.

Additionally, cyclic compression tests were conducted for 20,000 cycles at a frequency of 1 Hz, with hydrogels compressed up to 50% strain. The collected data were used to develop stress-strain curves and calculate mechanical parameters including recovery (i.e., the ratio of the area under loading curves in two specific cycles), energy dissipation (i.e., the difference between area under the curve of loading and unloading), and dissipation percentage. All hydrogels were equilibrated in PBS at 37 °C for 24 hours prior to testing.

### 2.7. Differentiation of hiPSCs to VICs and VECs

The use of human cells in this study was approved by the Ethical Committee of CHU Sainte-Justine Research Center (2025-8135). The cellular response to the fabricated hydrogels was assessed using valvular interstitial cells (VICs) and valvular endothelial cells (VECs) derived from human induced pluripotent stem cells (hiPSCs). The protocol for preparing hiPSC-derived VICs and VECs was adapted from a previously published method ^[16]^. Briefly, hiPSCs were expanded in mTeSR1 medium (Stemcell Technologies, Canada) until reaching the desired confluency. To start the differentiation, hiPSCs were detached and reseeded at a density of approximately 4 × 10^4^ cells/cm² in Matrigel-coated plates. After incubation for 24 hours in mTeSR1 medium, the media was replaced with a 1:1 mixture of DMEM/F12 (Gibco, USA) and Neurobasal medium (Gibco, USA), supplemented with 8 μM of CHIR-99021 (Selleck, USA), 25 ng/mL BMP4 (Gibco, USA), 1.94% B27 (Gibco, USA), and 0.97% N2 (Gibco, USA). The cells were cultured in this medium for 3 days. On day 4, this media was switched to StemPro-34 SFM (Gibco, USA) supplemented with 1 μM forskolin (Selleck, USA) and 200 ng/mL VEGF-165 (Gibco, USA). Differentiation was continued for an additional 3 days, with the media being changed daily. To confirm the success of the differentiation, at the end of day 6, cells were characterized by flow cytometry and immunofluorescent staining.

#### 2.7.1. Flow cytometry

Differentiated cells were detached and dissociated into single cells by TrypLE™ express enzyme (Gibco, USA). Live cells surface antigen were blocked with 5% fetal bovine (FBS, Gibco, USA) in PBS. Endothelial surface markers CD144 and CD31 were stained by incubating the cells with Alexa Fluor 647 mouse anti-human CD144 and FITC Mouse Anti-Human CD31 (BD Biosciences, USA), at 1/25 dilution for 30 min. All steps were carried out on ice. Flow cytometry data were acquired using a FACSCanto II (BD Biosciences) and analyzed by Flowjo software (Tree Star).

#### 2.7.2. Immunofluorescent staining

For staining, cells were fixed using 4% formaldehyde solution for 10 min at room temperature. Fixed cells were then permeabilized using tween-20 (Sigma Aldrich, USA), followed by blocking with 1 % bovine serum albumin (BSA, Sigma Aldrich, USA) solution in PBS supplemented with 5% goat serum (Gibco, USA). Subsequently, cells were incubated with CD 144 (Alexa Fluor 647 mouse anti-human CD144), CD 31 (BD Biosciences, USA), or alpha-smooth muscle actin (α-SMA, cy-3 conjugated α-SMA, Sigma Aldrich, USA) antibodies overnight at 4 °C. Nuclei were stained with DAPI (Invitrogen, USA) for 10 mins. Antibody dilutions were prepared according to the manufacturer’s recommendations. Stained cells were imaged using a fluorescent microscope (DMi-8, Leica Microsystems).

### 2.8. In vitro cell response

Following differentiation, VICs and VECs were dissociated using TrypLE™, counted, and seeded onto 3D printed hydrogels at a density of 2.5 × 10⁵ cells per printed hydrogel. Hydrogels with a diameter of 10 mm were 3D printed for this experiment. The cell-coated hydrogels were cultured in a 1:1 mixture of endothelial growth medium (EGM-2, Lonza, Switzerland), supplemented with 2 µM forskolin, and high glucose Dulbecco’s Modified Eagle Medium (DMEM, Gibco, USA) supplemented with 10% FBS (Gibco, USA). At day 4, 7, and 14, samples were removed from the incubator and stained using a Live/Dead viability kit (Invitrogen, USA) following the manufacturer’s instructions. For immunofluorescent staining, samples were stained for CD144, α-SMA, or CD31 as described above. Images were captured using a TCS SP8 confocal microscope (Leica Microsystems). Cells cultured on standard cell culture plates served as the control and were imaged using a fluorescent microscope (DMi-8, Leica Microsystems).

### 2.9. In vitro Hydrodynamic Testing

The in vitro functionality of the 3D printed heart valves was evaluated using a pulse duplicator system (Vivitro Labs, Canada). Tri-leaflet heart valves were 3D printed using the formulated inks and the DLP 3D printer. Following fabrication, the valves were swollen in water for 24 hours. Valves were then mounted on supports and positioned in the ventricular outlet of the pulse duplicator. A 23 mm Edwards trileaflet valve was placed in the tricuspid position at the atrial outlet. Testing was conducted under pulmonary conditions with a stroke volume of 70 mL and a heart rate of 70 beats per minute (BPM). Ventricular and pulmonary pressures were recorded using fiber optic pressure sensors (FISO Technologies, Canada). To assess leaflet coaptation and valve dynamics during opening and closing, ultrasound imaging and optical video recordings were performed. The performance of the printed heart valves was compared to a biological heart valve of similar size (Edwards Lifesciences), which served as a control. Water was used as the testing fluid in the experiments. The maximum opening area percentage was calculated using the recorded videos and represents the maximum passing area opening compared to the valve’s inlet.

### 2.10. Subcutaneous implantation

All animal procedures were conducted in accordance with the regulations of the Canadian Council on Animal Care (CCAC) and were reviewed and approved by the Ethical Committee of Centre hospitalier de l’Université de Montréal (CHUM, 2025-12661, C24042MLr). Twelve male Sprague-Dawley rats (180-200g, Charles River Laboratories) were used for this study. Animals were randomly divided to four groups of control, week 4, week 8, and week 12 and individually housed and allowed to acclimate for a minimum of 48 hours before any procedures, with ad libitum access to food and water. Environmental enrichment was provided throughout their stay. On the surgery day, animals received a subcutaneous injection of Buprenorphine SR (1 mg/kg) for pre-emptive analgesia 3 hours before surgery. Anesthesia was induced with 2-3% isoflurane in an induction chamber and maintained with 1-2% isoflurane via a facial mask for the duration of the procedure. The dorsal surface of the rat was shaved and aseptically prepared using alternating scrubs of chlorhexidine and alcohol. Four full-thickness dermal incisions were made, and subcutaneous pockets were created to implant the hydrogels (disks of 1 cm in diameter). The incisions were then closed using an intradermal suture pattern with 4-0 Vicryl (a resorbable suture). Buprenorphine SR provided analgesia for approximately 72 hours post-operatively. At the pre-determined endpoints (4, 8, and 12 weeks, n=3 per group), animals were euthanized via an intraperitoneal overdose of sodium pentobarbital. The hydrogel implants and surrounding tissue were then excised for subsequent analysis. For bulk RNA seq controls, subcutaneous pockets were made, but no hydrogel was implanted. The tissue surrounding the incision was retrieved and used as control.

### 2.11. Histological and immunohistochemistry staining

For histological analysis, explanted hydrogels and the surrounding host tissues were immediately fixed in 10% neutral-buffered formalin (VWR, Canada) following retrieval. After fixation, specimens were processed through graded ethanol dehydration, cleared, and embedded in paraffin. Paraffin blocks were sectioned at 7 µm thickness using a rotary microtome (RM2145, Leica). Sections were stained according to the manufacturer’s protocols using the following kits: Hematoxylin and Eosin (H&E; AB245880, Abcam, USA), Masson’s Trichrome (AB150686, Abcam, USA), Picro Sirius Red (AB150681, Abcam, USA), Von Kossa (AB150687, Abcam, USA), and Movat’s Pentachrome (AB245884, Abcam, USA).

For immunohistochemistry (IHC), freshly harvested tissues were embedded in optimal cutting temperature (O.C.T.) compound (Tissue-Plus™, Fisher Scientific, Canada), rapidly frozen on dry ice, and stored at −80 °C until further processing. Cryosections were prepared using a cryostat (CryoStar™ NX50, Thermo Scientific, USA) and mounted onto glass slides. Sections were fixed in 4% paraformaldehyde, permeabilized, and blocked prior to antibody incubation. Primary antibodies included CD3 (14-0030-82, Thermo Fisher Scientific, Canada), CD45 (AB10558, Abcam, USA), CD68 (AB125212, Abcam, USA), CD86 (942-MSM1-P0, Thermo Fisher Scientific, Canada), CD163 (MCA342GA, Bio-Rad, USA), vimentin (AB92547, Abcam, USA), and α-smooth muscle actin (α-SMA; AB7817, Abcam, USA). Sections were subsequently counterstained using Alexa Fluor™ 488 (A-11001, Thermo Fisher Scientific, Canada) and Alexa Fluor™ 647 (A-31573, Thermo Fisher Scientific, Canada) and mounted using mounting media.

All stained sections were imaged using a slide-scanning microscope (Axioscan, Zeiss). Digital images were analyzed by expert pathologists to assess host tissue response to the implanted hydrogels. For IHC staining, images were analyzed using QuPath 6.0.6 to calculate the positive area per DAPI, a total of 4 fields per sample, and minimum 3 samples were used.

### 2.12. Bulk RNA sequencing

Approximately 10mg of tissue surrounding the subcutaneously implanted biomaterial was excised and immediately frozen in N_2_ vapor. For long-term storage, samples were kept at-80 °C. RNA from these samples was extracted using ReliaPrep™ RNA Miniprep Systems/Tissues/50 preps (Promega, Z6111) according to manufacturer protocol. RNA was measured by NanoDrop 2000 spectrophotometer (Thermo Fisher Scientific). The RNA quality was further checked with Agilent Bionaylizer and RNA with RNA Integrity Number (RIN) above 7 were used for sequencing The DRAGEN RNA v4.3.16 pipeline was used for the analysis. Raw FASTQ reads were aligned to the Rattus norvegicus (rn6, Ensembl 110) genome with the DRAGEN RNA spliced aligner, allowing for read soft-clipping to exclude low-quality and adapter sub-sequences. The DRAGEN pipeline quantifies gene and transcript expression using an expectation-maximization algorithm to infer the best transcript assignment for each read.

Bulk RNA sequencing data were analyzed using the DESeq2 package in R. Genes with a total read count of ≤10 across all samples were excluded to remove lowly expressed genes. The filtered count matrix was used as input for DESeq2, and sample metadata were provided to define the experimental conditions (Control, Week 4, Week 8, and Week 12), with three biological replicates per group. Differential gene expression analysis was performed using the DESeq2 pipeline. Size factors were estimated to account for differences in sequencing depth between samples, followed by model fitting using the negative binomial distribution. P values were adjusted for multiple testing using the Benjamini–Hochberg procedure to control the false discovery rate (FDR). Regularized log (rlog) transformation was applied to the count data for downstream visualization. Principal component analysis (PCA) was performed to assess sample distribution and variability across experimental conditions. Heatmaps were generated using the pheatmap package. A predefined panel of genes associated with inflammatory responses, immune-cell markers, complement pathways, cytokine signaling, and innate immune responses was investigated. For this targeted analysis, normalized expression values were standardized across samples for each gene using row-wise Z-score transformation. Samples were organized according to their experimental groups. Volcano plots were generated using the EnhancedVolcano package to visualize differential gene expression. Log2 fold changes were plotted against adjusted P values, with thresholds of |log2FC| > 2 and an adjusted P value below 0.01 used to highlight differentially expressed genes.

Gene-level counts were imported into R, and downstream transcriptomic analyses were performed including principal component analysis (PCA) to assess global transcriptional variation, heatmap visualization of gene-expression patterns, and pathway enrichment analysis to identify biological processes and signaling pathways associated with the observed transcriptional changes.

### 2.13. Statistical analysis

All experiments were conducted in triplicate unless stated otherwise. Data are represented as the mean ± standard deviation (SD). Statistical analysis was performed to identify significant differences using t-test or ANOVA where applicable, with a p-value of less than 0.05 considered statistically significant. A total of 4 different significance was used: *p < 0.05, **p < 0.01, ***p < 0.001.

## 3. Results and discussion

### 3.1. Rationale for GPQuin formulation

DLP printing is a light-based 3D printing technique that has gained significant attention in recent years for the fabrication of TESs. DLP offers several advantages over other 3D printing methods, such as eliminating the high shear stresses, higher printing resolution against extrusion-based bioprinting, and achieving faster printing speeds compared to stereolithography (SLA) printing ^[12, 17]^. Nevertheless, the application of DLP printing for tissue engineering applications is currently hindered by the scarcity of suitable photocurable inks ^[18]^. To date, materials such as GelMA, PEGDA, silk fibroin, and chitosan have been explored for DLP 3D printing ^[17, 19]^. GPQuin was formulated by combining photocurable synthetic and naturally derived polymers with complementary intended roles. Low-molecular-weight PEGDA (i.e., Mw 700 Da) was used to provide a high concentration of reactive alkene groups, enabling rapid cross-linking and enhanced printing resolution. However, at high concentrations low-molecular-weight PEGDA hydrogels are mechanically brittle. A mixture of low- and high-molecular weight PEGDA was used to reduce the necessity for using high concentrations of low-M_w_ PEGDA. Moreover, PEGDA is bioinert and lacks cell adhesion sites, which is a key drawback for its applications where cell attachment and structure remodeling over time is required ^[20]^. Consequently, GelMA was incorporated into GPQuin as a photocurable gelatin derivative containing cell-interactive motifs. Furthermore, GelMA is photosensitive, making it a suitable candidate for DLP printing. Finally, QSGM was included as a photocrosslinkable polysaccharide component previously developed by our group^[14]^. Its polymer backbone photosensitivity and pendant functional groups provide a basis for possible additional chemical and physical interactions within the composite, e.g., hydrogen bonding. These considerations guided the formulation design; however, the present study evaluates the resulting composite as a whole and does not independently resolve the contribution of each component or establish a specific intermolecular reinforcement mechanism.

### 3.2. 3D DLP printing of GPQuin

When combined with an appropriate photoinitiator and light source, polymerization and crosslinking of the photosensitive ink can be effectively initiated. The DLP instrument used in our experiments was equipped with a 405 nm visible light projector. Accordingly, LAP was selected as the photoinitiator. LAP has a molar extinction coefficient of approximately 25 M⁻¹cm⁻¹ at 405 nm, making it particularly well-suited for biofabrication applications that utilize visible light for photocrosslinking. In addition, LAP is highly water-soluble, which facilitates its use in aqueous bioink formulations, allowing the fabrication of cell-laden hydrogel scaffolds ^[21]^. Apart from the ink composition, the success of printing complex constructs with high resolution and fine details is also highly dependent on the printing conditions, including light source intensity, exposure time, and curing layer thickness. These parameters directly influence the degree of crosslinking and, consequently, affect the printing fidelity and resolution ^[22]^. Printing conditions were selected using a test structure containing features of different dimensions (**Figure 2a**). For GPQuin ink, the selected settings were a layer thickness of 100 µm, a projector power setting of 50%, a base-layer exposure of 40 s, and a body-layer exposure of 10 s. These settings were used for the constructs described below (**Figure 2a**). It is important to note that these parameters are sensitive to changes in ink formulation or layer thickness. For example, reducing the layer thickness would typically require a shorter exposure time due to decreased light penetration depth.

**Figure 2.**
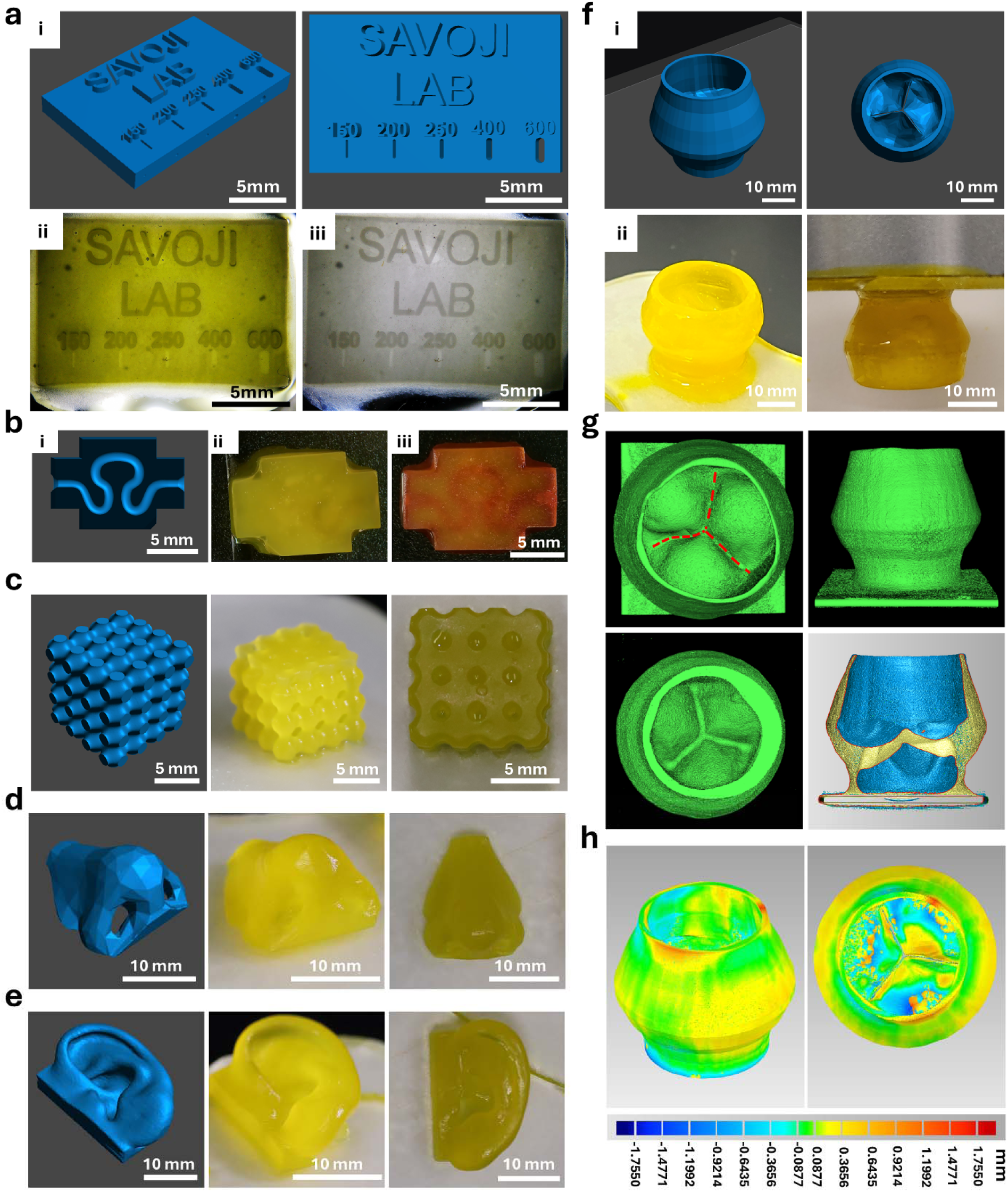
GPQuin can print a variety of geometries. (a) Evaluating the printing fidelity: (i) STL design used to characterize the printing fidelity, and optical images showing the printed structures with optimum printing parameters post printing (ii) and after 24 h in PBS (iii). (b) GPQuin was used to print a perfusable scaffold: (i) STL design, (ii) the printed scaffold before perfusion, and (iii) after perfusion with colored water. The design and post-printing optical images of the DLP-printed TESs: (c) A cube with Schwarz P volumetric lattice, (d) an anatomically analogous nasal structure, and (e) an ear with comparable anatomical features. (f) 3D printing of TEHVs using GPQuin: (i) The STL design used for printing and (ii) optical images of the TEHVs after print. (g) µCT scan renders of the 3D printed TEHVs post-printing. The dashed red lines show the leaflet coaptation location and the plane cuts of the µCT renders showing the presence of leaflets and sinus of Valsalva within the 3D printed TEHVs. (h) Surface-deviation maps overlaying of the µCT reconstruction with the corresponding STL model.

The selected printing conditions enabled fabrication of various geometries (**Figure 2b-e**). First, GPQuin was used to print scaffolds with internal channel. Upon the printing, the presence of a continuous hollow channel was confirmed by the perfusion of the channel with colored water (**Figure 2b** and supplementary video S1). To further validate the ink’s versatility, a cube featuring a Schwarz Primitive was printed (**Figure 2c**). The printed cube successfully retained the micron-scale features of the original design, as evidenced by the presence of well-defined hollow spaces within the geometry. Additionally, anatomically analogous models of a human nose (**Figure 2d**) and ear (**Figure 2e**) were successfully printed to showcase the potential of GPQuin for producing geometrically complex, human-scale constructs. Optical imaging revealed high fidelity of the printed structures as compared to the initial CAD design. These examples illustrate the range of geometries accessible with the formulation.

For the 3D printing of TEHVs GPQuin enabled fabrication of 25 mm trileaflet valve scaffolds with separated leaflets and anatomically inspired sinuses (Figure 2f-h). Early macroscopic investigations after printing revealed that the 3D printed TEHVs closely mimic the STL design (Supplementary video S2). Micro-computed tomography (µCT) confirmed preservation of tissue-specific features, including the valve leaflets and sinus of Valsalva. Comparison of the reconstructed surface with the reference STL yielded mean positive and negative deviations of +214 and −248 µm, respectively. Compared to our previous attempts using extrusion-based bioprinting to fabricate the same design (+0.52 overprinting and -0.50 mm underprinting)^[11]^, this valve exhibited almost two-folds improved structural printing fidelity. This enhancement can be attributed to the inherently higher resolution capabilities of DLP printing, which allows for more precise creation of fine features.

### 3.3. Physicochemical characteristics of the DLP-printed Hydrogels

The swelling and degradation behavior of the 3D printed hydrogels were investigated as two key physicochemical properties of the fabricated constructs. The high water affinity inherent to hydrogels enables them to more closely mimic the native ECM, which is critical for replicating physiological conditions. Moreover, the swelling capacity of hydrogels plays a vital role in facilitating the diffusion of nutrients and oxygen toward the inner parts of the hydrogel, thereby enhancing cell infiltration and survival ^[23]^. 3D printed hydrogels showed a high swelling of 877.3 ± 14.2 %, showing their hydrophilic nature (**Figure 3a**). To compare, hydrogels from 4% GelMA were printed using the same printing parameters, and the swelling percentages were found to be 1863.3 ± 50.4 %. GPQuin exhibited lower swelling than the 4% GelMA reference under the tested conditions. Since the formulations differ in both composition and total polymer content, this comparison does not isolate the effects of PEGDA or QSGM. The observed difference is consistent with differences in network composition and structure, although crosslink density was not directly measured.

**Figure 3.**
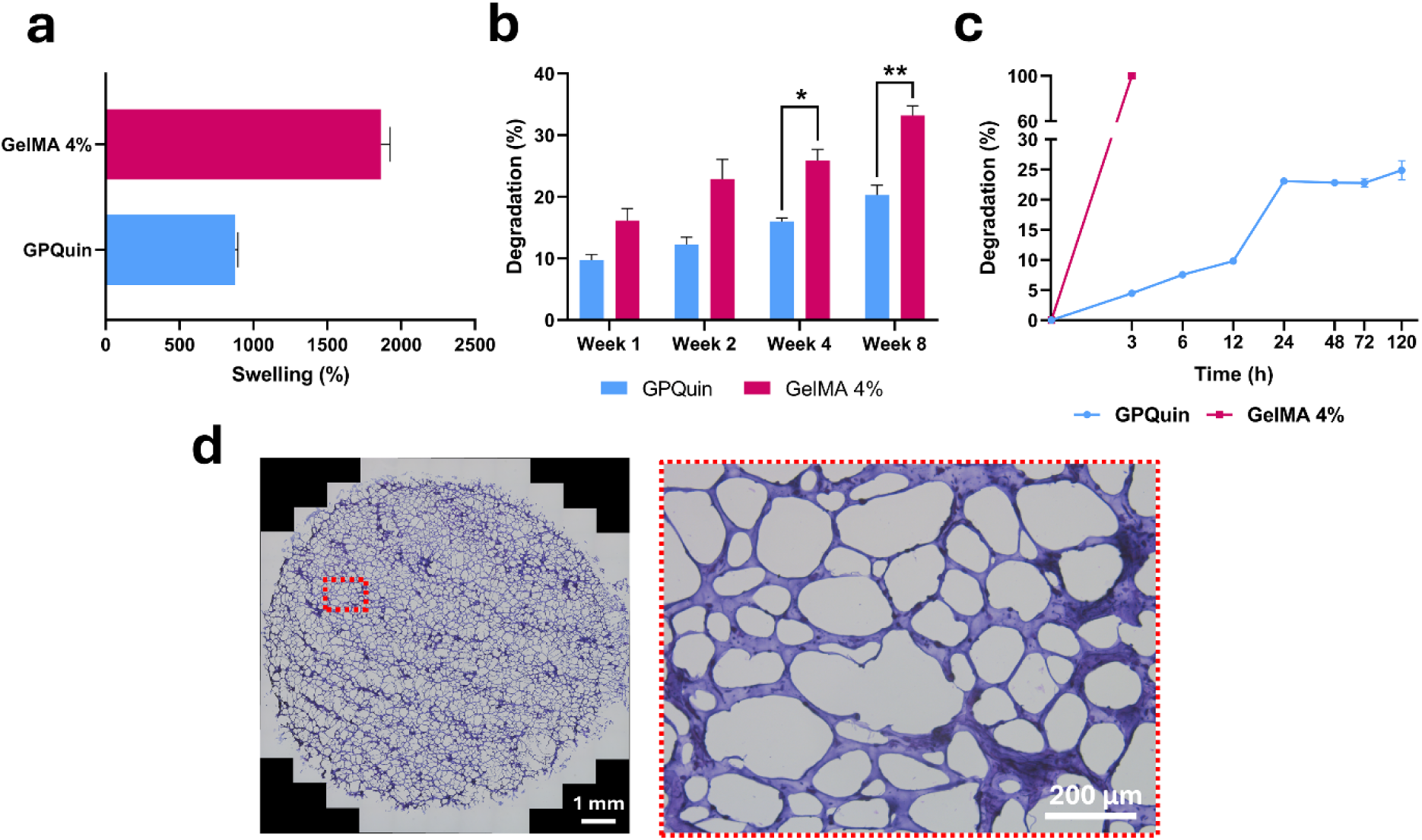
Physicochemical and structural characterization of the DLP-printed GPQuin hydrogels. (a) Equilibrium swelling, (b) hydrolytic degradation, and (c) enzymatic degradation of the formulated hydrogels. 3D printed GelMA hydrogels (4 wt.%) were used as control. (*p<0.05, **p <0.01). (d) Representative H&E-stained cryosection of 3D-printed hydrogels illustrating the internal structure.

The hydrolytic and enzymatic degradation of the GPQuin hydrogels was also recorded and compared with GelMA 4% hydrogels. As depicted in **Figure 3b, 3D** printed hydrogels showed 9.7 ± 0.8, 12.2 ± 1.0, 16.0 ± 0.5, and 20.27 ± 1.62 % degradation percentages after 1, 2, 4, and 8 weeks, respectively. On the other hand, the degradation percentage for GelMA alone hydrogels was found to be 16.2 ± 1.6, 22.9 ± 2.6, 26.9 ± 1.6, and 33.20 ± 1.58 % at the same time points. This significant reduction in degradation rate, similar to the swelling, could be attributed to differences in composition, polymer content, and network structure. Higher polymer concentration could result in increased chemical bonding as well as physical interactions, which in turn reduce the degradation speed of the hydrogels and enhance their stability.

The enzymatic degradation profiles of the 3D printed hydrogels and 4% GelMA scaffolds were also assessed under enzymatic digestion. Under collagenase exposure, the 4% GelMA hydrogel reached complete measured mass loss within 3 h, whereas GPQuin exhibited approximately 25% mass loss after 120 h (**Figure 3c**). The GPQuin therefore retained a larger proportion of its initial mass under these conditions. This is consistent with greater structural stability of the composite network, although, as noted above, this comparison does not isolate the contributions of PEGDA or QSGM. These findings are consistent with previously reported trends in similar hydrogel systems, where GelMA-based scaffolds exhibit relatively rapid degradation, while composite formulations demonstrate enhanced structural stability ^[24]^.

The microstructure of the 3D printed hydrogels was investigated through cryosectioning and H&E staining (**Figure 3d**). An apparent continuous microporous architecture was observed across the entire cross-section of the printed hydrogels, which likely accounts for their high swelling capacity of the hydrogels. Furthermore, this structural feature suggests that the hydrogels can facilitate adequate perfusion of oxygen and nutrients to infiltrating cells over time. Nevertheless, careful consideration must be given to pore size, as the measured dimensions may be artificially changed due to ice crystal formation during the freezing process and sectioning process which imposes stress on the hydrogels.

### 3.4. Mechanical characteristics of GPQuin Hydrogels

The mechanical microenvironment of heart valves is unique, and valve leaflets experience a variety of mechanical stresses, including bending, compressive, shear, and tensile stresses ^[25]^. These stresses are cyclic in nature, and valves should experience and withstand such stresses millions of times during their lifetime. Consequently, it is critically important to evaluate the mechanical properties of the developed scaffolds to ensure their capability of tolerating such a dynamic microenvironment. To this end, the mechanical performance of the 3D printed hydrogels was assessed through tensile and compressive tests (**Figure 4** and supplementary Figure S1). The tensile properties of the hydrogels are presented in Figure 4b and c and supplementary Figure S1. Based on the obtained results, the hydrogels showed an ultimate tensile strength (UTS) of 113.2 ± 17.4 kPa with an elongation at break of 75.3 ± 2.3 %. Moreover, Young’s modulus of the hydrogels in the early stage of elongation (<20%) was found to be 117.2 ± 15.5 kPa. When compared to native adult human tissue, the modulus of the developed hydrogels was in the same order of magnitude as the low, pretransitional region along the circumferential direction of the posterior mitral leaflet (PML, 180.1 ± 71.5 kPa),tricuspid valve leaflets (TV, 102.6 ± 14.4 kPa) reported in the literature^[26]^. Moreover, the increase in modulus from 117.2 ± 15.5 kPa below 20% strain to 191.3 ± 32.0 kPa between 20% and 50% strain indicates strain-dependent stiffening within the tested range.

**Figure 4.**
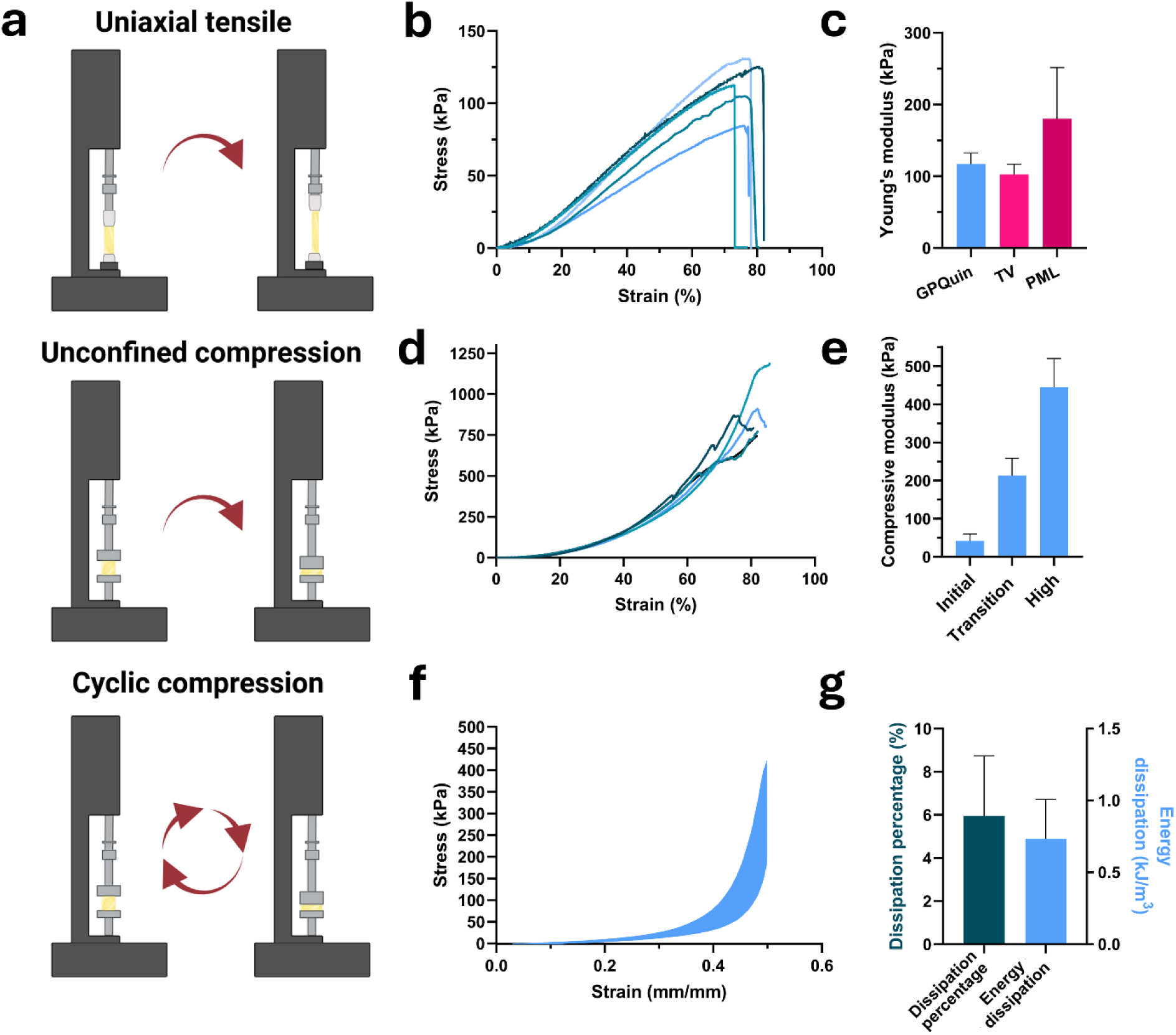
Mechanical characterization of DLP-printed GPQuin hydrogels. (a) Schematic depiction of performed mechanical tests. (b) The tensile strain-stress curves recorded during the uniaxial tensile tests (n=5). (c) Calculated Young’s modulus of the hydrogels. Represented data for PML and TV are obtained from Ref. ^[26]^. (d) Compressive curves (n=5) recorded during the compressive tests with (e) their subsequent calculated compressive modulus. (f) Stress–strain curves of cycles 2 to 10,000 from cyclic compression test for GPQuin hydrogel and (g) calculated energy dissipation and dissipation percentage.

The compressive mechanical properties of the hydrogels were investigated through an unconfined uniaxial compression test (**Figure 4d** and **e**). As expected, the 3D printed hydrogels exhibited a range of stiffness, from 42.0 to 444.5 kPa, based on the compression percentage, indicating non-linear and dynamic changes in the mechanical properties of the hydrogels. Furthermore, the hydrogels showed high compressive resilience, with compressive strains around 80%. These observations can be attributed to the extensive chemical and physical interactions between various components of the hydrogels, resulting in a robust hydrogel.

Finally, to assess mechanical stability under repetitive loading, the GPQuin hydrogels were subjected to 20,000 compression cycles at 1 Hz and up to 50% strain (**Figure 4f and 4g, and Supplementary Figure S1**). No macroscopic rupture was observed throughout the test, indicating suitable structural integrity under cyclic loading. The hydrogels exhibited an average energy dissipation of 0.73 ± 0.28 kJ m⁻³ per cycle, corresponding to an energy dissipation ratio of 5.9 ± 2.8%. The cyclic response evolved during loading, with smaller relative changes between later cycles. Recovery values were 42.4%, 61.9%, and 85.1% when comparing cycle 20,000 with cycles 2, 2,000, and 10,000, respectively. This suggests that the cyclic loading response starts to stabilize after the initial loading cycles, with only reduced changes observed thereafter, suggesting capability of sustained resistance to repeated compression. This behaviour is likely attributable to the combined chemical crosslinking and physical interactions within the hydrogel network, which together provide a mechanically robust structure.

### 3.5. Characterization of hiPSC-derived cells and their response to GPQuin hydrogels

hiPSCs were differentiated toward VECs and VICs using a two-stage protocol, originally reported by Cai et al. ^[16]^. Based on their proposed mechanism, hiPSCs were initially directed toward cardiac lateral plate mesoderm linage by activation of BMP canonical Wnt signaling pathways. This is then followed by utilizing of VEGF and activation of NFATc1 signaling pathway to promote VECs, as described previously (**Figure 5a**). Following differentiation, the resulting cells were analyzed using the flow cytometry technique (**Figure 5b**). CD31 and CD144 were selected as the general endothelial surface markers expressed by VECs to identify and distinguish them from the other cell populations. Differentiation efficiency was assessed by flow cytometry, using CD31 and CD144 as endothelial markers (**Figure 5b**). A total of 69.3 % of the cells were both CD31^+^ and CD144^+^, indicating that the majority of the population had successfully differentiated into endothelial cells. When compared to the original report, we closely replicated differentiation efficiency (70.8%) of the previous work. Immunofluorescence analysis further validated these findings, with the majority of cells co-expressing CD31 and CD144 (**Figure 5c**). In parallel, α-SMA-positive cells were detected in cells negative for endothelial markers, confirming the presence of a subpopulation of VICs within the differentiated cells. Together, these results demonstrate efficient differentiation of hiPSCs into mixed valvular cell populations suitable for subsequent cell–material interaction studies. However, it should be noted that these results do not independently establish that the differentiated cells possess mature valve-specific identity.

**Figure 5.**
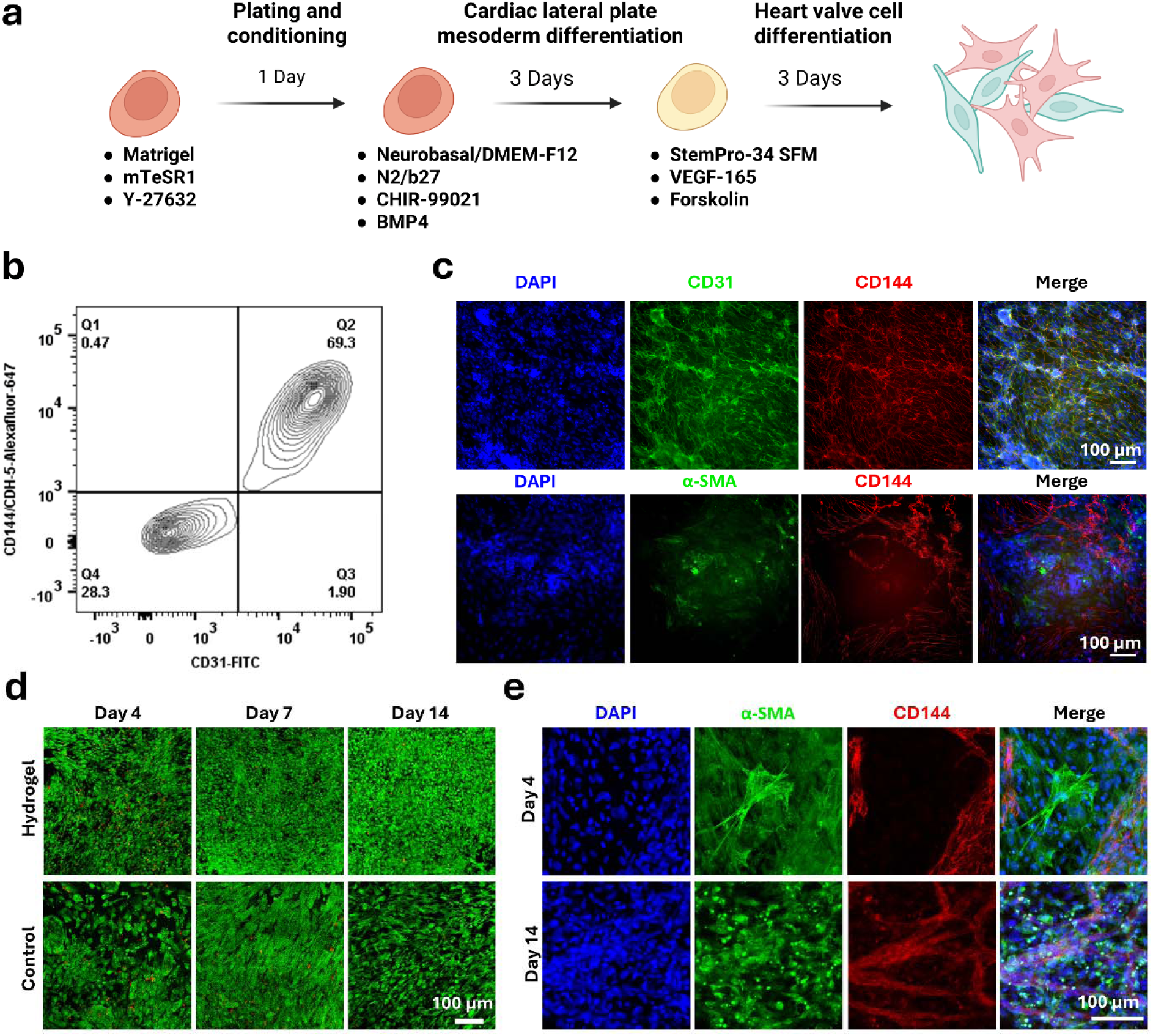
Differentiation of hiPSCs into valve-specific cell types and biological assessment of the GPQuin hydrogels. (a) Schematic illustration of the stepwise differentiation protocol for VECs and VICs. (b) Flow cytometry analysis of differentiated cells. (c) Immunofluorescent staining of differentiated cells for endothelial markers CD31 and CD144 and α-SMA, showing the presence of both VECs and VICs in the cell population, Scale bar 100 µm. (d) Live/Dead staining of hiPSC-derived VICs and VECs cultured on the 3D printed hydrogels at days 4, 7, and 14. Live cells appear green, and dead cells appear red. Scale bar 100 µm. (e) Immunofluorescent staining of hiPSC-derived VICs and VECs cultured on hydrogels for DAPI (nuclei), α-SMA (VIC marker), and CD144 (VEC marker), Scale bar 100 µm.

We next evaluated the cytocompatibility of GPQuin hydrogels using hiPSC-derived VECs and VICs. High biocompatibility and low cytotoxicity are essential characteristics of TESs to support effective recellularization and integration over time. Differentiated cells were directly seeded onto DLP-printed GPQuin hydrogels, and viability was evaluated after 4, 7, and 14 days. using Live/Dead fluorescent staining (**Figure 5d**). At all time points, cells remained attached to the hydrogels and exhibited high viability, with the majority of cells stained green, indicating minimal cytotoxicity. Furthermore, cells preserved their phenotype over the two weeks of culture. No significant visual differences in cell viability were observed between the hydrogel group and the control group (cell culture plates), demonstrating the suitability of the hydrogels for supporting the survival of valve-specific cell populations.

Further characterization of cell phenotype and behavior on the hydrogels was performed using cell-specific immunofluorescent staining (**Figure 5e**). Similar to the differentiation step, CD144 was used to identify VECs, while α-SMA served as the marker for VICs ^[16]^. On day 4 post-seeding, both CD144^+^ and α-SMA^+^ cells were observed on the hydrogel surface, confirming the successful attachment and coexistence of both cell types. Morphologically, α-SMA^+^ cells exhibited an elongated, polygonal shape characteristic of VICs, whereas CD144^+^ cells displayed a cobblestone-like morphology, typical of endothelial cells. Similar staining patterns and cellular phenotypes were maintained through day 14, suggesting that both hiPSC-derived VECs and VICs preserved their lineage-specific phenotypes over time. These results indicate favorable cell-material interactions and further support the biocompatibility of the hydrogel scaffold for 14-day culture of valve-like cell types.

### 3.6. In vivo biocompatibility assessment of the GPQuin Hydrogels

#### 3.6.1. Histological and IHC evaluations

The in vivo biocompatibility of the GPQuin hydrogel was investigated through a 12-week subcutaneous implantation follow up (Figure 6a). The explanted hydrogels and surrounding tissues were subjected to comprehensive histological analysis using H&E, Masson’s Trichrome, Picrosirius Red, von Kossa, and Movat’s pentachrome staining to evaluate the host tissue response (Figure 6b and Supplementary Figure S2). Based on the observations, the implanted biomaterial was consistently surrounded by a well-defined fibrous capsule, in keeping with a typical chronic foreign body response. Hydrogel fragments were observed extending beyond the implant core into the surrounding capsule and, in some cases, into the adjacent tissue with a maximal migration distance of around 750 μm. Interestingly, the greatest extent of hydrogel dissemination was observed at 8 weeks post-implantation. No evidence of intravascular embolization of biomaterial particles was identified in the examined sections. The implant–tissue interface demonstrated a localized inflammatory response associated with variable degrees of fibrosis and neovascularization arising within the developing fibrous capsule. A granulomatous inflammatory reaction characterized by multinucleated foreign body giant cells was consistently observed surrounding extravasated hydrogel fragments.

**Figure 6.**
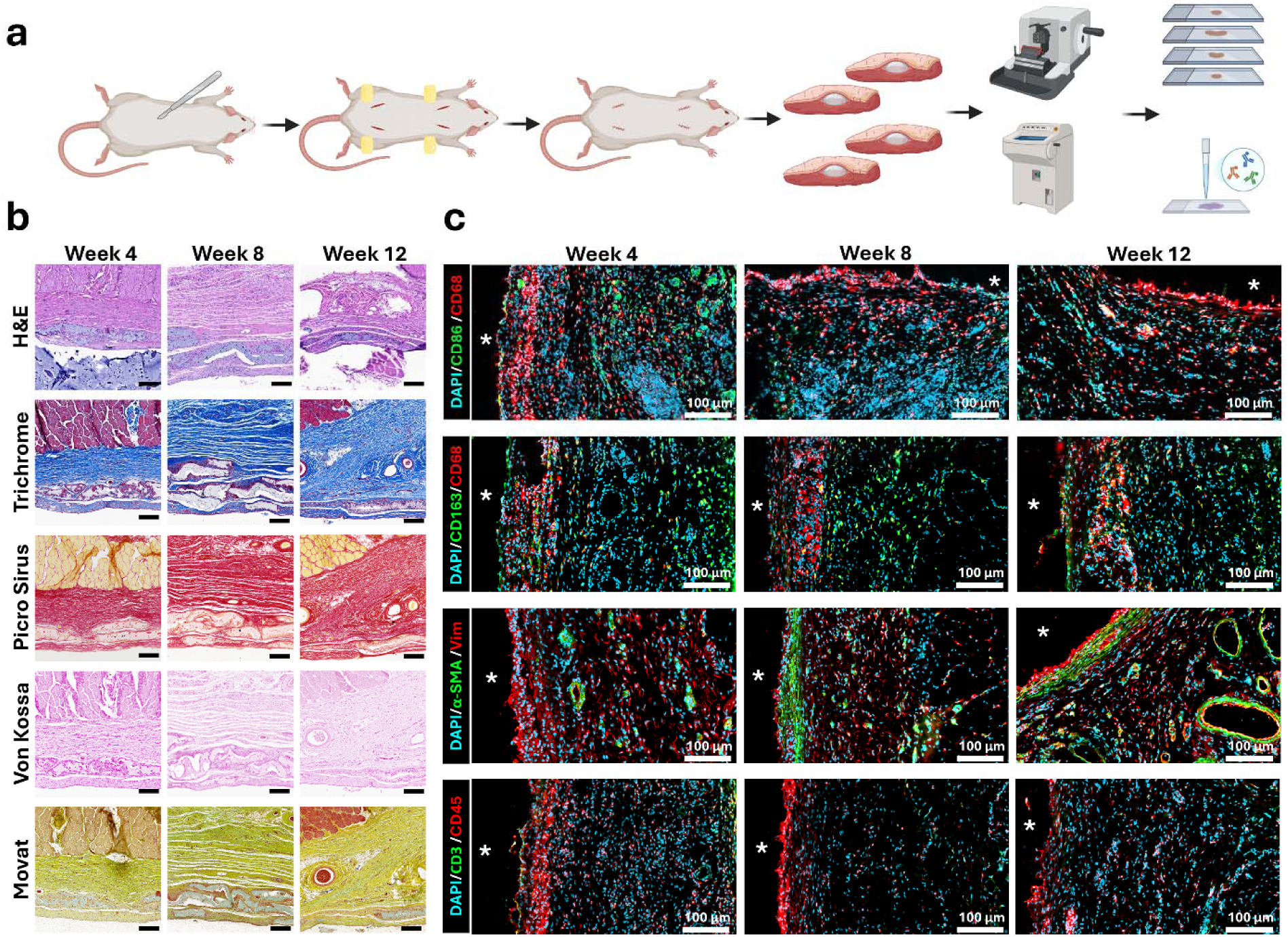
In vivo biocompatibility assessment of 3D printed GPQuin hydrogels using a rat subcutaneous model. (a) Schematic representation of the performed surgical procedure for the test. (b) Histological staining images of explanted hydrogels to assess general response, inflammation, ECM deposition, and absence of calcification of the hydrogels during the implantation. Scale bar = 250 µm. (c) Fluorescent IHC staining of the explanted scaffolds using selected biomarkers (CD3, CD45, CD68, CD86, CD163, Vimentin, and α-SMA). Scale bar = 100 µm. White asterisk (*) shows the location of the hydrogel in the vicinity of the tissue.

The inflammatory response showed a gradual decline over time, with the highest cellularity at week 4 and a reduction continuing through weeks 8 and 12. The reduction was characterized by decreased numbers of lymphocytes, plasma cells, macrophages, and giant cells. In contrast, the thickness and organization of the fibrous capsule appeared to increase over time. Neovascularization was more prominent at week 4 and appeared to either stabilize or mildly regress by week 12. No significant polymorphonuclear (neutrophilic) infiltrate was identified at any time point. Additionally, there was no evidence of tissue necrosis or adipose tissue infiltration in the examined sections. Overall, the examined sections showed reduced inflammatory cellularity at later time points together with persistent, increasingly organized fibrous encapsulation.

We further investigated inflammatory cell infiltration around the subcutaneous implants by immunostaining (**Figure 6C and Supplementary Figure S4-8**). Immunostaining for the common monocyte/macrophage marker CD68, together with CD86 and CD163 as markers of inflammatory and anti-inflammatory macrophages, respectively, revealed a higher number of CD68^+^ cells at early time point, which decreased over the subsequent 8 weeks and was accompanied by an increase in CD163^+^ cells (**Figure 6C, Supplementary Figure S4a and b and Supplementary Figure S5 and 6**). These observations may indicate a partial shift from a pro-inflammatory toward an anti-inflammatory macrophage profile over time. We also assessed CD45^+^ cells as a general marker of both innate and adaptive immune cells. Similar to CD68^+^ cells, the number of CD45^+^ cells was lower at 12 weeks than at earlier time points (**Figure 6C, Supplementary Figure S4d, and Supplementary Figure S8**). In addition, assessment of CD45 in combination with CD3, a pan-T-cell marker, revealed less attenuation of the cellular inflammatory response. This finding is consistent with the RNA-seq data presented in the next section and the general trend of a slower adaptive response relative to the innate immune response.

We also assessed fibrosis at the cellular level by measuring α-SMA expression, which tends to increase in fibroblasts, endothelial cells, and SMCs during fibrosis. Enhanced α-SMA expression in Vim^+^ cells from 8 weeks onward (**Figure 6C and Supplementary Figure S4c**) suggests that fibrosis begins after the acute inflammatory phase and tends to surround and stabilize around the biomaterial.

#### 3.6.2. Bulk RNA-seq

To further assess the body’s reaction to GPQuin, upon hydrogel explantation the RNA was extracted, and bulk RNA sequencing on poly-A-enriched samples was performed (**Figure 7**). Principal component analysis visualized group-associated variation in gene expression (**Figure 7a)**. GO biological pathway analysis (**Figure 7d-f**) showed that inflammatory pathways involved in leukocyte recruitment and activation, along with collagen biosynthesis pathways, were among the most upregulated at 4 weeks vs. control. After 8 weeks, cellular metabolism changed relative to 4 weeks, and cells became more metabolically active; inflammatory pathways remained active but shifted toward antigen processing and presentation, with the emergence of adaptive immunity pathways compared with controls. At 12 weeks after implantation, GPQuin samples showed some metabolic shift relative to earlier time points, but pathways involved in collagen deposition and ECM remodeling remained active relative to sham-operated control tisssue. Adaptive immune pathways in B lymphocytes were significantly more active at later time points, suggesting memory formation against the biomaterial and the development of active humoral immunity.

**Figure 7.**
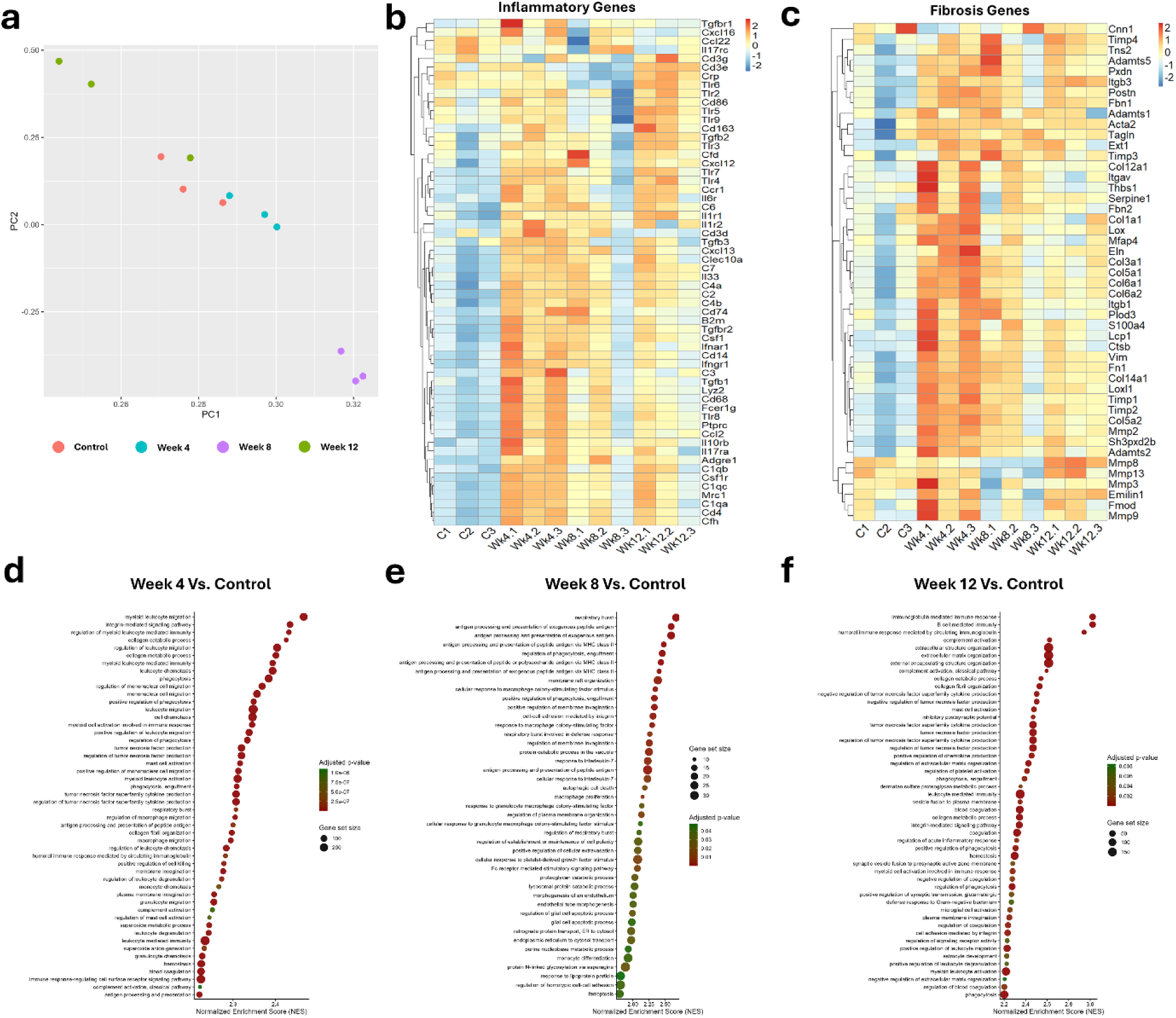
Transcriptomic profiling of the body’s response to GPQuin scaffolds. (a) Principal component analysis (PCA) plot illustrating the gene expression variation across experimental groups. Heatmaps displaying z-score normalized gene expression profiles for key markers associated with (b) inflammatory pathways and (c) ECM remodeling and fibrosis across all experimental groups. Gene Set Enrichment Analysis (GSEA) showing top enriched Gene Ontology (GO) biological processes relative to Control at (d) Week 4, (e) Week 8, and (f) Week 12 post-implantation.

A closer look at the gene expression changes underlying the altered pathways showed that GPQuin hydrogels led to a temporary increase in inflammatory cells from both innate and adaptive immune responses (**Figure 7b**). At week 4 post-implantation, general macrophage-dendritic cell markers, which are important players in innate immune response, including CD68 and CD14 ^[27]^, were increased, but their presence attenuated over the next 8 weeks. In contrast, the increase in T-lymphocyte markers (CD3, CD4, and CD8) was lower in magnitude but more persistent, maintaining stable levels of expression following biomaterial implantation, a trend indicative of adaptive immunity. In addition to the presence of these markers, early post-implantation up-regulation of Toll-like Receptor (TLR) family genes and complement pathway components indicated that the biomaterial induced macrophage recruitment via elevated adhesion molecule expression (**Figure 7 and Supplementary Figure S9**). These recruited macrophages recognize and phagocytose the biomaterial using innate immune systems for foreign object and microbial detection. Over time, expression of these markers, along with nuclear factors and key regulators of inflammatory pathways, gradually declined (**Supplementary Figure S9**), signaling the resolution of inflammation and a transition toward a pro-healing macrophage phenotype. Overall, while early CD14 expression and TLR activation mediated initial foreign body sensing and clearance, the concurrent involvement of T cells highlights a coordinated adaptive immune response ^[28]^.

Alongside inflammation, the degree of fibrosis serves as a critical parameter for evaluating biomaterial biocompatibility, typically manifesting as ECM deposition driven by an imbalance between ECM synthesis and degradation ^[29]^. Genes associated with matrix production and remodeling also showed time-dependent expression patterns. At 4 weeks post-implantation, ECM deposition marked by elevated collagen and elastin expression increased, accompanied by upregulation of myofibroblast differentiation markers (**Figure 7c, and Supplementary Figure S9**). Upstream signaling regulators—including TGF-β, Smad3, and Snail2—alongside key matrix remodeling enzymes such as matrix metalloproteinases (MMPs) and a disintegrin and metalloproteinases (ADAMs),^[30]^ were similarly upregulated. By week 8, expression levels of these profibrotic genes attenuated. However, at week 12, matrix remodeling genes like MMPs exhibited a secondary, modest increase ^[29]^. Given the slow turnover of ECM at the transcriptional level, this late upregulation suggests that established fibrotic tissue undergoes sustained, active remodeling by week 12, consistent with ongoing remodeling of the fibrous capsule. Together, these data indicate a foreign-body response in which inflammation declines over time while fibrous encapsulation and matrix modeling persists. This controlled inflammatory and fibrotic sequence which is characterized by transient activation rather than progressive scarring, underscores the favorable biocompatibility of the biomaterial ^[31]^. However, since the analysis was performed on bulk tissue, the contributions of changing cell abundance and altered expression within individual cells cannot be separated at this stage.

### 3.7. In vitro hydrodynamic characteristics

Proper function under physiological conditions is the most important characteristic of a TEHV. Thus, we evaluated the functionality of DLP-printed GPQuin TEHVs under physiological pulmonary conditions using a pulse duplicator system operating at a heart rate of 70 bpm and a stroke volume of 70 mL (**Figure 8a**). Overall, the valves remained structurally intact during testing, with no visible signs of tearing or macroscopic damage (**Figure 8b**). We further benchmarked their performance against a commercially available bioprosthetic valve with a comparable inflow diameter. Video analysis throughout the cardiac cycle showed leaflet opening and apparent coaptation during closure without prolapse in both valve types (**Figure 8c, and Supplementary Video S3**), indicating that the GPQuin TEHVs maintained functionality under pulmonary loading. Analysis of leaflet motion using echocardiograhy further demonstrated compliant leaflet deformation and apparentcoaptation during closure, consistent with appropriate valve function (**Supplementary Video S4**). However, the bioprosthetic valve exhibited a larger maximum opening area (58% vs 24%). The smaller opening of the GPQuin valve may reflect differences in leaflet geometry, thickness, material properties, or mounting.

**Figure 8.**
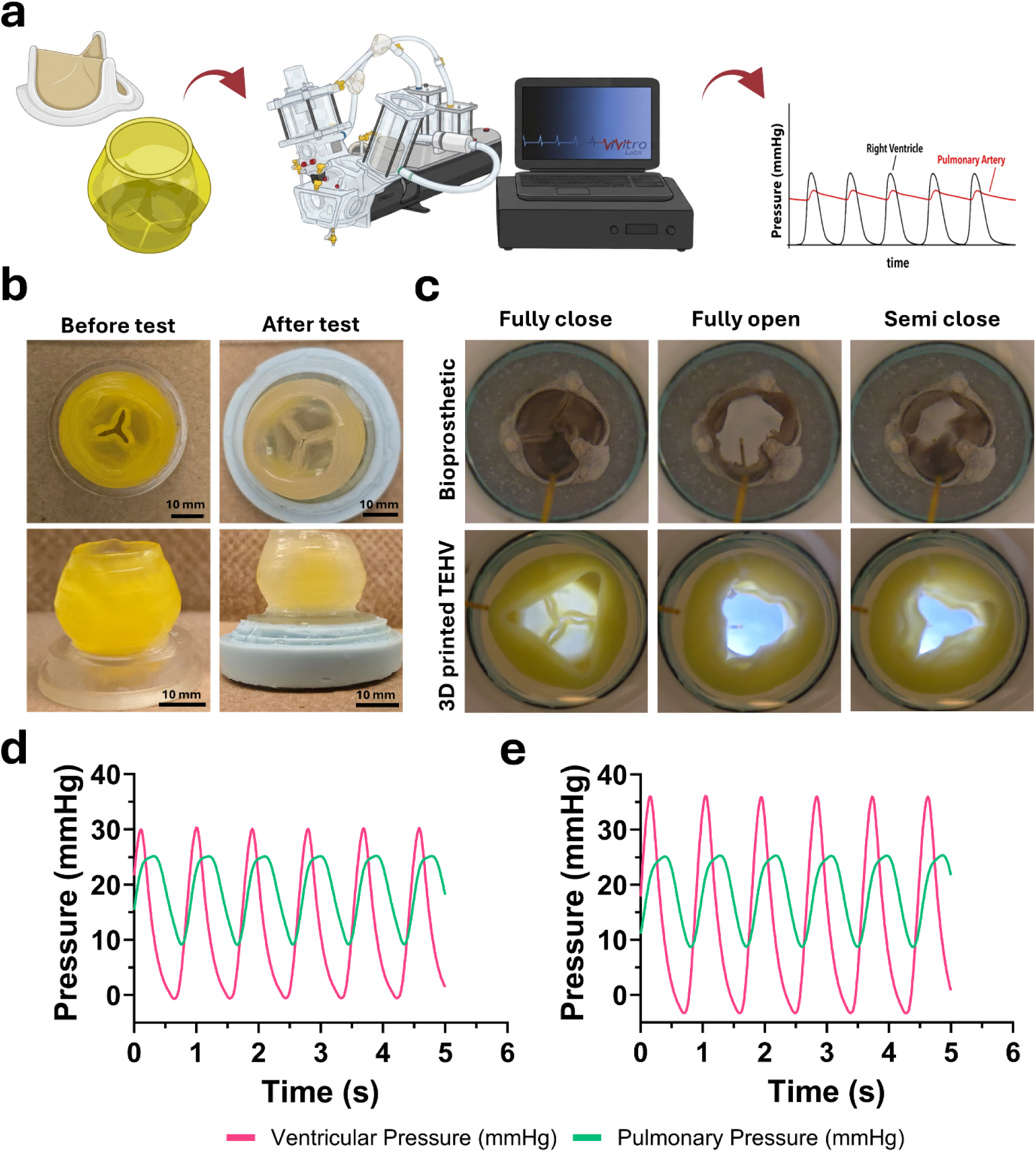
Evaluation of the DLP-printed GPQuin TEHVs under pulsatile flow. (a) Schematic representation of in vitro hemodynamic assessment setup. (b) Macroscopic optical images of a TEHV before (left) and after (right) performing hydrodynamic assessment under physiological conditions. Scale bar = 10 mm. The leaflet shortening before the test is due to dehydration. (c) Representative optical images showing the bioprosthetic and TEHV valves at different stages of a cardiac cycle. Pressure versus time graphs recording the ventricular and pulmonary pressures downstream and upstream of the (d) bioprosthetic valve and (e) the DLP-printed GPQuin TEHV.

We next quantified the hemodynamic performance of the valves by recording upstream and downstream transvalvular pressures under pulmonary conditions (**Figure 8d and e**). Both valves generated physiological pressure waveforms throughout the cardiac cycle. The bioprosthetic valve exhibited a peak-to-peak pressure gradient of 4.8 mmHg, determined directly from pressure measurements obtained using transducers positioned immediately upstream and downstream of the valve in the pulse duplicator , which likely resulted from a mismatch between the valve size and the pulse duplicator outflow tract. The DLP-printed GPQuin TEHVs exhibited a higher pressure gradient of 10.4 mmHg, consistent with the smaller maximum opening area observed during video analysis and the valve geometry. Despite this increase, the valves maintained stable opening and closure dynamics throughout testing, supporting their functionality under pulmonary valve conditions and their suitability for subsequent in vivo evaluation.

## 4. Conclusions and Outlook

In conclusion, this study presents a novel multi-material bioink, GPQuin, which combines PEGDA, GelMA, and methacrylated quince seed mucilage in a photocurable formulation for digital light processing (DLP) 3D printing, enabling the fabrication of complex, human-sized scaffolds with high precision Our results underscore the advantages of DLP 3D printing over traditional extrusion-based methods, particularly in terms of print fidelity and the ability to incorporate complex geometries and micron-sized features within the constructs. The developed hydrogels, demonstrated mechanical properties close to native heart valve tissues and showed suitable biocompatibility and cytocompatibility, supporting the culture of valve-specific cell types derived from hiPSCs. Furthermore, the demonstrated in vitro functionality and potential for in vivo remodeling of the 3D printed GPQuin hydrogels highlighted their potential for the development of TESs. Finally, the observed performance outcomes of the DLP printed GPQuin heart valve under physiological hemodynamic conditions, exhibited their functionality potential to be used as a TEHV. As such, this work lays a strong foundation for future studies exploring the in vivo translation of DLP-printed TEHVs, with the aim of providing a viable solution for patients requiring valve replacement.

## Author Contributions

**Arman Jafari**: conceptualization, methodology, investigation, formal Analysis, visualization, validation, writing – original draft, writing – review and editing, data curation. **Alireza Nemati**: investigation, formal Analysis, visualization, writing – review and editing. **Arman Sarshoghi**: methodology, investigation, formal Analysis, software, visualization, writing – review and editing. Naimeh Rafatian: investigation, validation, writing – original draft, writing – review and editing. **Sandrine Guenette**: formal Analysis, writing – original draft, writing – review and editing. **Yeojin Park**: investigation, writing – review and editing. **Vahid Karamzadeh**: methodology, writing – review and editing. **Ali Hedayatnia**: methodology, writing – review and editing. **Charles Leduc**: resources, writing – review and editing. **Moishe Liberman**: resources, writing – review and editing. **Ismail Bouhout**: resources, writing – review and editing. **Lyes Kadem**: resources, investigation, writing – review and editing. **Gregor Andelfinger**: resources, writing – review and editing. **Houman Savoji**: Supervision, funding acquisition, validation, project administration, resources, writing – review and editing.

## Acknowledgment

Our work is supported by the Natural Sciences and Engineering Research Council of Canada (NSERC; RGPIN-2021-03960 and DGECR-2021-00337), New Frontiers in Research Fund (NFRF), NSERC-FRQ NOVA program for junior researchers, Stem Cell Network, Fonds de Recherche du Québec Santé (FRQS) (Chercheurs-boursiers J1 (313837) and Chercheurs-boursiers J2) ), Establishment of Young Investigators (324277), Montreal TransMedTech Institute (iTMT), Start-up funds from the CRCHU Sainte-Justine (CRCHUSJ), University of Montreal, and Polytechnique Montreal to H.S

A.J. gratefully acknowledges the Fonds de Recherche du Québec - Santé (FRQS) Scholarship. We also acknowledge Mohammad Reza Habibi-Kavashkohie for his assistance with the analysis of bulk RNA-sequencing data.

## Data availability

Data will be made available on request.

## References

[1] H. Bauser-Heaton, O. M. Barry, S. C. Hofferberth, J. T. Tretter, M. Ma, A. Goldstone, A. Armstrong, T. K. Jones, A. Yoganathan, P. del Nido, JACC: Advances 2024, 3, 101191.

[2] a)C. Xu, K. Yang, Y. Xu, X. Meng, Y. Zhou, Y. Xu, X. Li, W. Qiao, J. Shi, D. Zhang, Journal of Nanobiotechnology 2024, 22, 378; b)Q. Tong, J. Cai, Z. Wang, Y. Sun, X. Liang, Q. Xu, O. A. Mahamoud, Y. Qian, Z. Qian, Small 2024, 20, 2309844; c)P. Zilla, P. Human, T. Pennel, Frontiers in Cardiovascular Medicine 2024, 11, 1347838.

[3] a)M. Y. Emmert, B. A. Schmitt, S. Loerakker, B. Sanders, H. Spriestersbach, E. S. Fioretta, L. Bruder, K. Brakmann, S. E. Motta, V. Lintas, Sci. Transl. Med. 2018, 10, eaan4587; b)J. Kluin, H. Talacua, A. I. Smits, M. Y. Emmert, M. C. Brugmans, E. S. Fioretta, P. E. Dijkman, S. H. Söntjens, R. Duijvelshoff, S. Dekker, Biomaterials 2017, 125, 101.

[4] R. Lanza, R. Langer, J. P. Vacanti, A. Atala, Principles of tissue engineering, Academic press, 2020.

[5] E. S. Fioretta, S. E. Motta, V. Lintas, S. Loerakker, K. K. Parker, F. P. Baaijens, V. Falk, S. P. Hoerstrup, M. Y. Emmert, Nature Reviews Cardiology 2021, 18, 92.

[6] J. Li, W. Qiao, Y. Liu, H. Lei, S. Wang, Y. Xu, Y. Zhou, S. Wen, Z. Yang, W. Wan, Nature Communications 2024, 15, 7462.

[7] a)D. F. V. Rey, J.-P. St-Pierre, in Handbook of Tissue Engineering Scaffolds: Volume One, Elsevier, 2019; b)A. D’Amore, S. K. Luketich, G. M. Raffa, S. Olia, G. Menallo, A. Mazzola, F. D’Accardi, T. Grunberg, X. Gu, M. Pilato, Biomaterials 2018, 150, 25; c)B. Mirani, S. O. Mathew, N. Latifi, M. R. Labrosse, B. G. Amsden, C. A. Simmons, Adv. Funct. Mater. 2024, 34, 2302786.

[8] A. Jafari, Z. Ajji, A. Mousavi, S. Naghieh, S. A. Bencherif, H. Savoji, Advanced Materials Technologies 2022, 7, 2101636.

[9] a)P. P. Borthakur, A. Das, J. J. Sahariah, P. Pramanik, E. Baruah, K. Pathak, Biomedical Materials & Devices 2025, 1; b)U. Ghosh, S. Ning, Y. Wang, Y. L. Kong, Adv. Healthc. Mater. 2018, 7, 1800417.

[10] a)A. Lee, A. Hudson, D. Shiwarski, J. Tashman, T. Hinton, S. Yerneni, J. Bliley, P. Campbell, A. Feinberg, Science 2019, 365, 482; b)M. B. Immohr, F. Dos Santos Adrego, H. L. Teichert, V. Schmidt, Y. Sugimura, S. Bauer, M. Barth, A. Lichtenberg, P. Akhyari, Biomedical Materials 2023, 18, 015004; c)L. Hockaday, K. Kang, N. Colangelo, P. Cheung, B. Duan, E. Malone, J. Wu, L. Girardi, L. Bonassar, H. Lipson, Biofabrication 2012, 4, 035005.

[11] A. Jafari, S. Vahid Niknezhad, M. Kaviani, W. Saleh, N. Wong, P. P. Van Vliet, C. Moraes, A. Ajji, L. Kadem, N. Azarpira, G. Andelfinger, H. Savoji, Adv. Funct. Mater. 2024, 34, 2305188.

[12] Y. Shen, H. Tang, X. Huang, R. Hang, X. Zhang, Y. Wang, X. Yao, Carbohydr. Polym. 2020, 235, 115970.

[13] a)L. Yue, S. Macrae Montgomery, X. Sun, L. Yu, Y. Song, T. Nomura, M. Tanaka, H. Jerry Qi, Nature Communications 2023, 14, 1251; b)H. Yang, M. Ji, M. Yang, M. Shi, Y. Pan, Y. Zhou, H. J. Qi, Z. Suo, J. Tang, Matter 2021, 4, 1935.

[14] A. Jafari, K. Bhatt, S. V. Niknezhad, A. Ajji, M. Griffith, G. Andelfinger, S. A. Bencherif, H. Savoji, Carbohydr. Polym. 2025, 352, 123140.

[15] a)A. Helling, E. Tsekoura, M. Biggs, Y. Bayon, A. Pandit, D. Zeugolis, ACS Biomater. Sci. Eng. 2017, 3, 1922; b)H. Yin, M. Zhu, Y. Wang, L. Luo, Q. Ye, B. H. Lee, Frontiers in Soft Matter 2023, 2, 1101680.

[16] Z. Cai, M. Zhu, L. Xu, Y. Wang, Y. Xu, W. Y. Yim, H. Cao, R. Guo, X. Qiu, X. He, Circulation 2024, 149, 1435.

[17] a)M. Zanon, R. Cue-López, E. Martínez-Campos, P. Bosch, D.-L. Versace, H. Hayek, N. Garino, C. F. Pirri, M. Sangermano, A. Chiappone, Additive Manufacturing 2023, 69, 103553; b)H. Hong, Y. B. Seo, J. S. Lee, Y. J. Lee, H. Lee, O. Ajiteru, M. T. Sultan, O. J. Lee, S. H. Kim, C. H. Park, Biomaterials 2020, 232, 119679.

[18] S. H. Kim, H. Hong, O. Ajiteru, M. T. Sultan, Y. J. Lee, J. S. Lee, O. J. Lee, H. Lee, H. S. Park, K. Y. Choi, Nat. Protoc. 2021, 16, 5484.

[19] M. Zanon, D. Baruffaldi, M. Sangermano, C. F. Pirri, F. Frascella, A. Chiappone, Eur. Polym. J. 2021, 160, 110813.

[20] H. G. Hosseinabadi, D. Nieto, A. Yousefinejad, H. Fattel, L. Ionov, A. K. Miri, Appl. Mater. Today 2023, 30, 101721.

[21] K. S. Lim, J. H. Galarraga, X. Cui, G. C. Lindberg, J. A. Burdick, T. B. Woodfield, Chem. Rev. 2020, 120, 10662.

[22] S. Swetha, T. J. Sahiti, G. S. Priya, K. Harshitha, A. Anil, Interactions 2024, 245, 178.

[23] a)D.-M. Radulescu, I. A. Neacsu, A.-M. Grumezescu, E. Andronescu, Polymers 2022, 14, 799; b)A. Jafari, A. Amirsadeghi, S. Hassanajili, N. Azarpira, Int. J. Pharm. 2020, 583, 119413.

[24] a)H. Yue, Y. Wang, S. Fernandes, C. Vyas, P. Bartolo, Macromol. Biosci. 2025, 25, 2400587; b)S. Das, R. Valoor, P. Ratnayake, B. Basu, ACS Appl. Bio Mater. 2024, 7, 2809.

[25] S. Arjunon, S. Rathan, H. Jo, A. P. Yoganathan, Ann. Biomed. Eng. 2013, 41, 1331.

[26] T. Pham, F. Sulejmani, E. Shin, D. Wang, W. Sun, Acta Biomater. 2017, 54, 345.

[27] a)D. A. Chistiakov, M. C. Killingsworth, V. A. Myasoedova, A. N. Orekhov, Y. V. Bobryshev, Lab. Invest. 2017, 97, 4; b)D. Sharygin, L. G. Koniaris, C. Wells, T. A. Zimmers, T. Hamidi, Immunology 2023, 169, 260.

[28] K. A. Fitzgerald, J. C. Kagan, Cell 2020, 180, 1044.

[29] A. E. Mayorca-Guiliani, D. J. Leeming, K. Henriksen, J. H. Mortensen, S. H. Nielsen, Q. M. Anstee, A. J. Sanyal, M. A. Karsdal, D. Schuppan, npj Metabolic Health and Disease 2025, 3, 25.

[30] X.-m. Meng, D. J. Nikolic-Paterson, H. Y. Lan, Nature Reviews Nephrology 2016, 12, 325.

[31] M. E. Dickenson, R. S. Oakes, A. H. Morris, npj Biomedical Innovations 2026, 3, 3.

