## Supplemental Materials for "Digital light processing of human-scale tissue-engineered heart valves"

* Corresponding author:

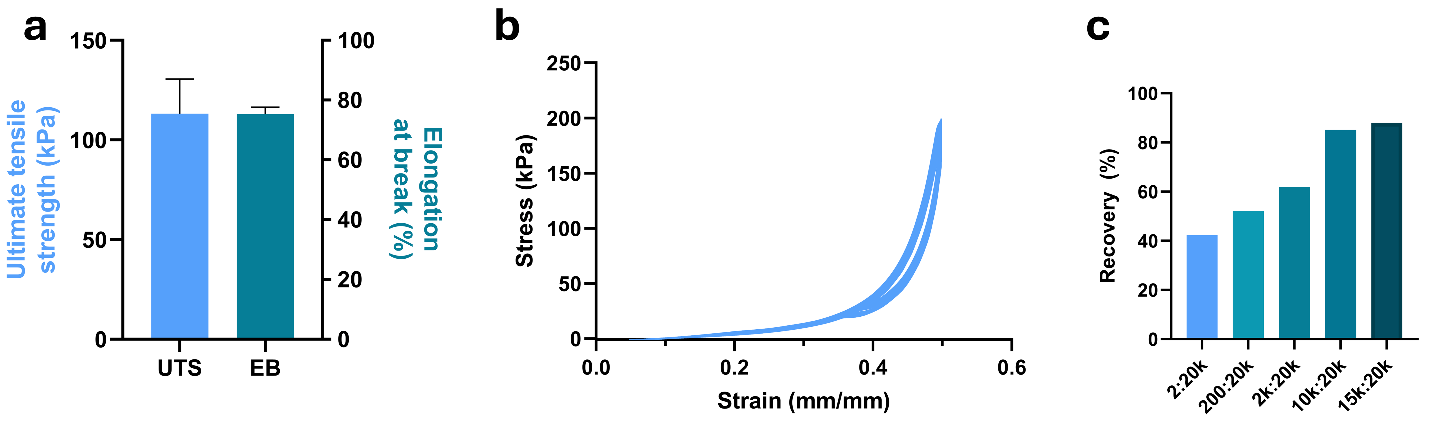

**Supplementary Figure S1**. (a) Calculated ultimate tensile strength (UTS) and elongation at break (EB) for the GPQuin hydrogels using tensile testing. (b) Stress-strain curves for cyclic compression between cycle 10,001 and 20,000 and (c) calculated recovery percentage between different cycle numbers.

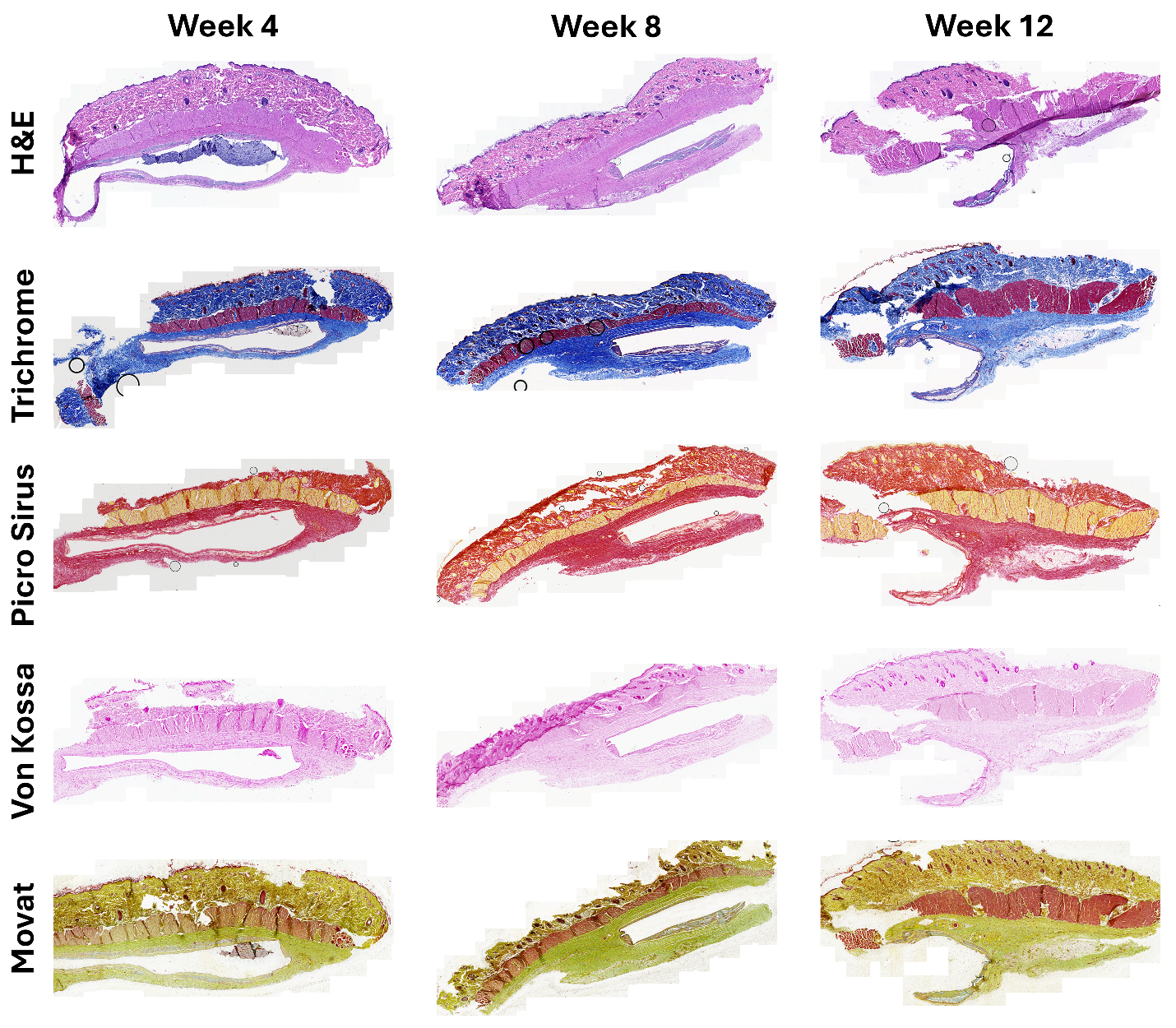

**Supplementary Figure S2.** Representative longitudinal sections from subcutaneous explants at weeks 4, 8, and 12 stained with H&E, trichrome, Picrosirus red, von Kossa, and Movat’s pentachrome.

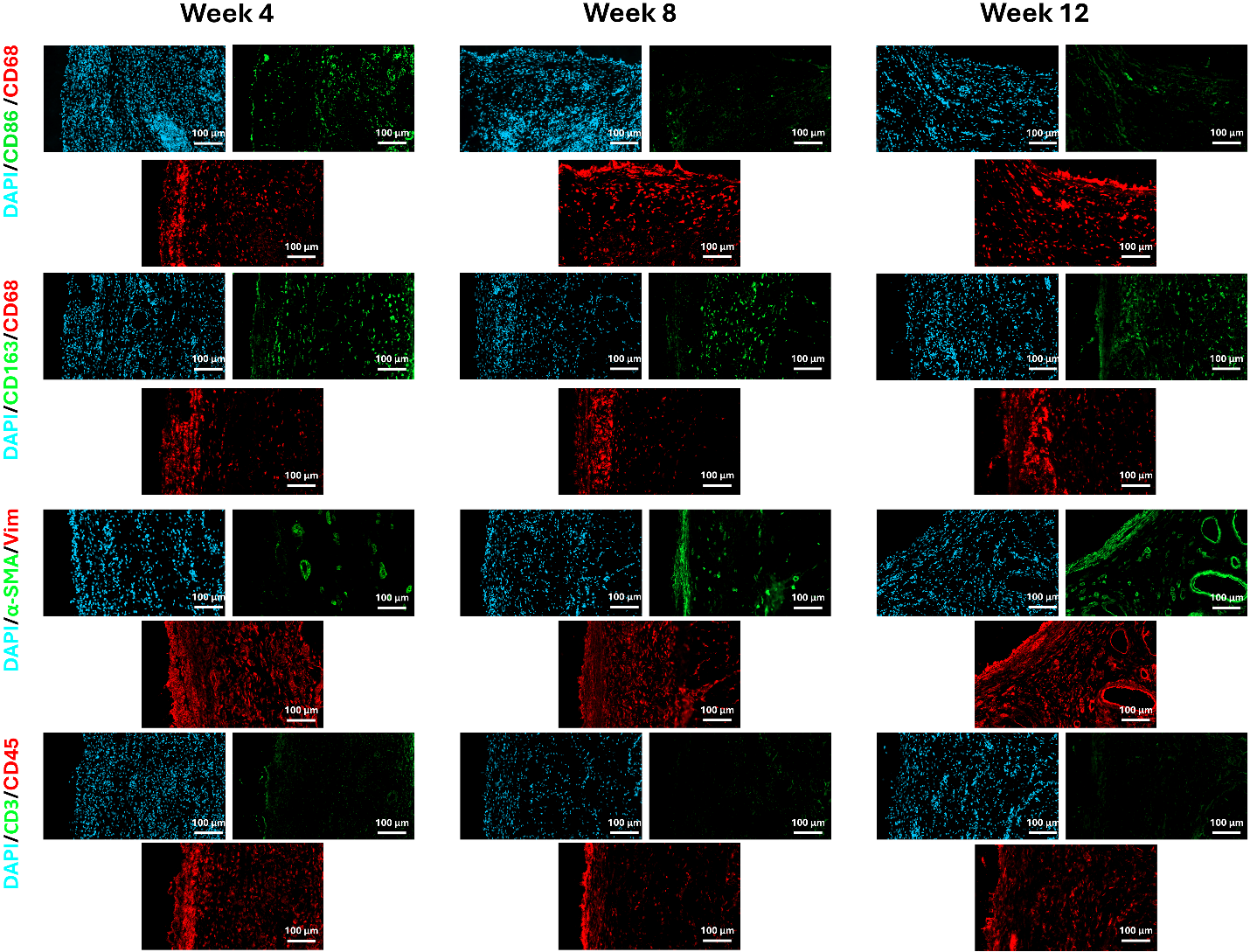

*****

*****

*****

*****

*****

*****

*****

*****

*****

*****

*****

*****

*****

*****

*****

*****

*****

*****

*****

*****

*****

*****

*****

*****

*****

*****

*****

*****

*****

*****

*****

*****

*****

*****

*****

*****

**Supplementary Figure S3.** Representative high magnification fluorescent IHC staining of the explanted scaffolds using selected biomarkers (CD3, CD45, CD68, CD86, CD163, Vimentin, and α-SMA). White asterisk (*) shows the location of the hydrogel in the vicinity of the tissue.

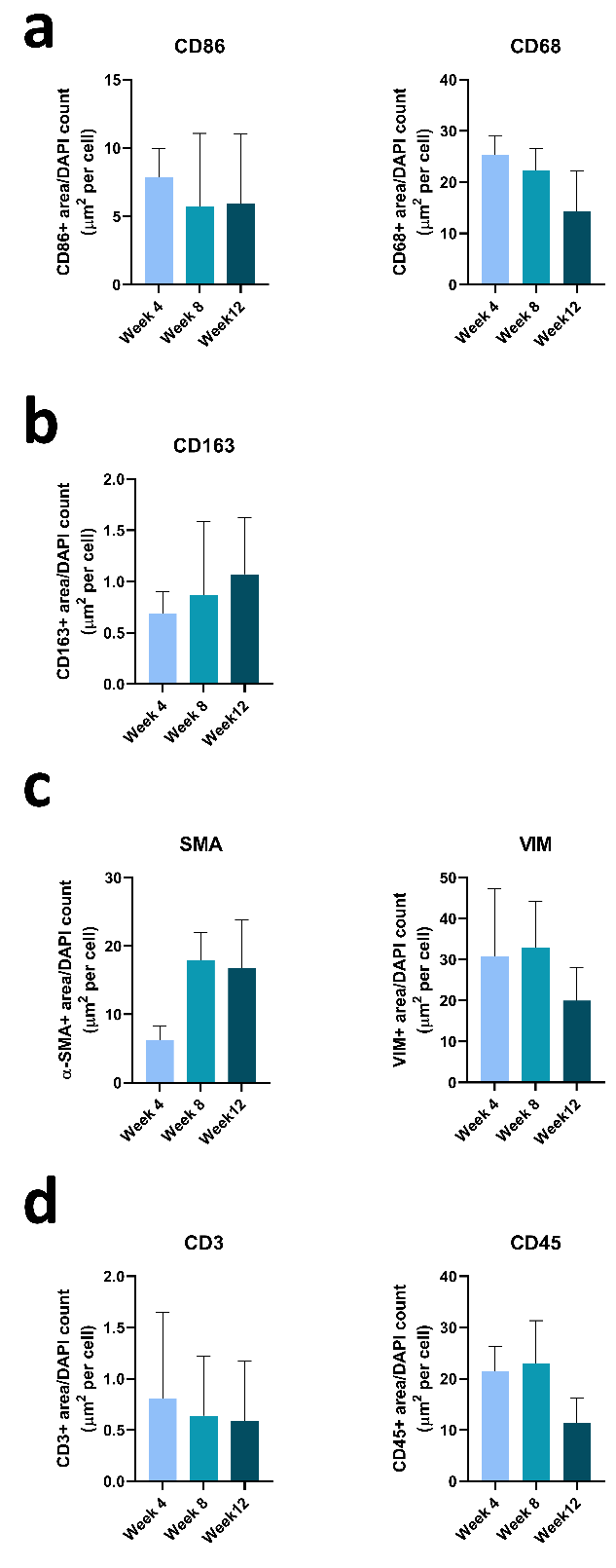

**Supplementary Figure S4.** Quantification of immunohistochemical staining for inflammatory, macrophage, stromal, and lymphocyte markers. (a) Quantification of CD86 and CD68, (b) CD163, (c) α-SMA and VIM, and (d) CD3 and CD45 markers at 4, 8, and 12 weeks. Staining was quantified as the marker-positive area normalized to the number of DAPI-positive cells.

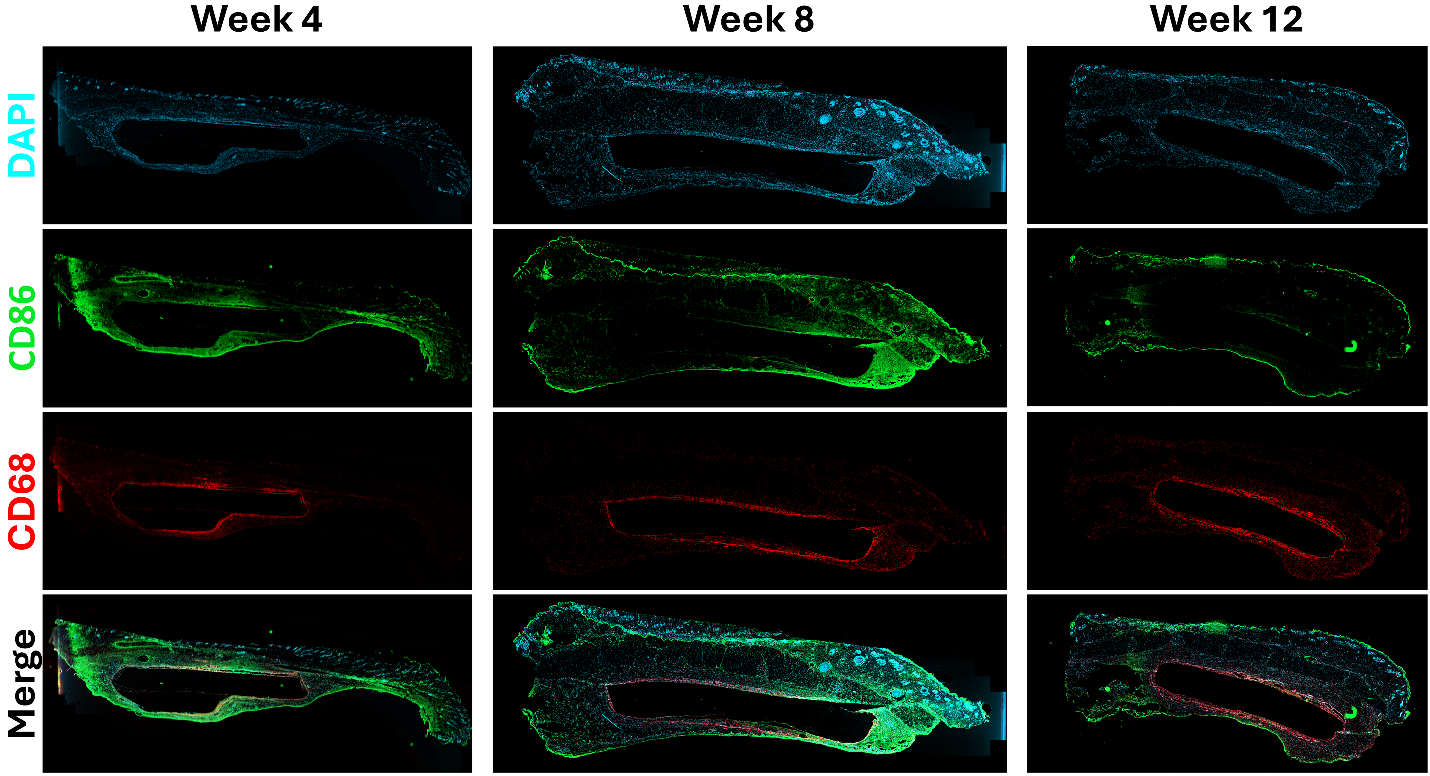

**Supplementary Figure S5.** Representative longitudinal sections from subcutaneous explants stained for DAPI (blue), CD86 (green), and CD68 (red).

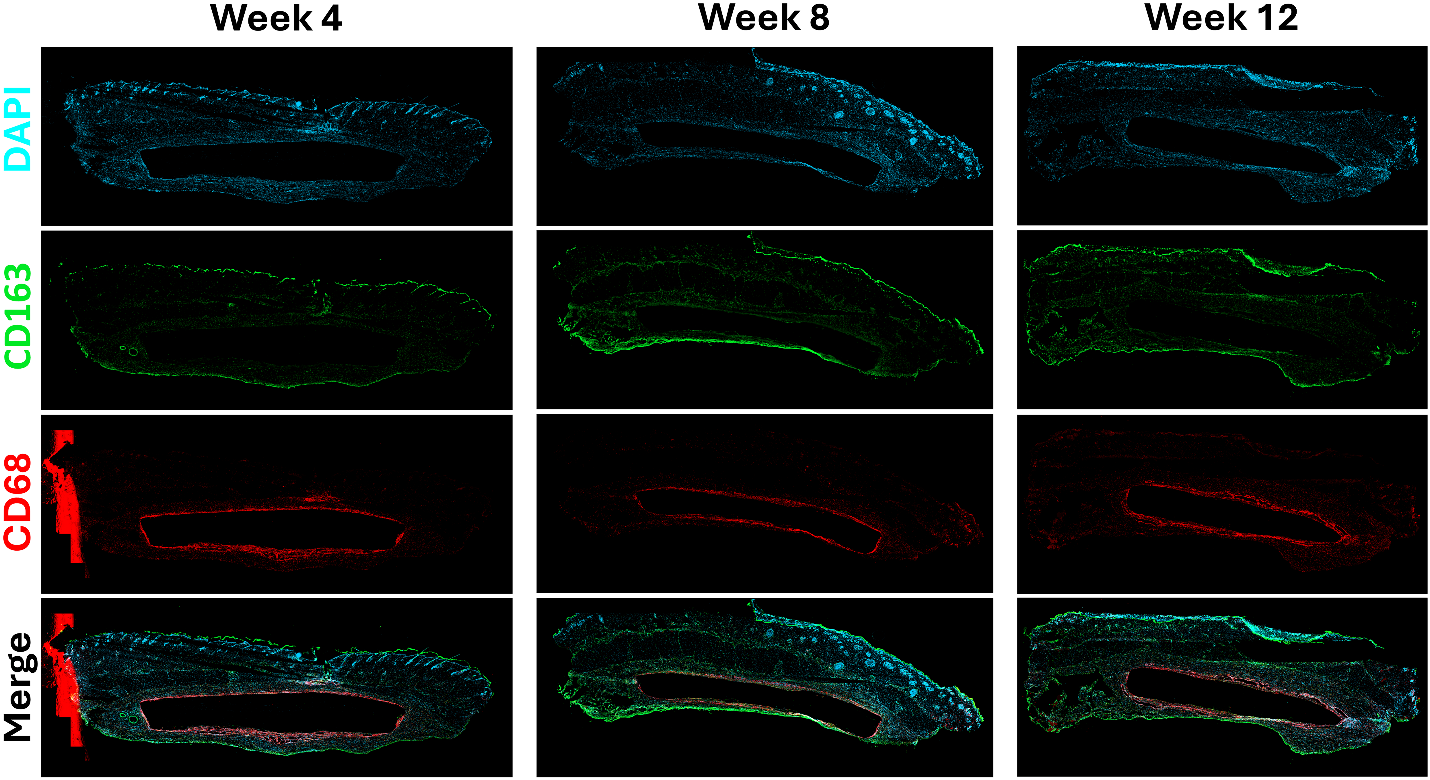

**Supplementary Figure S6.** Representative longitudinal sections from subcutaneous explants stained for DAPI (blue), CD163 (green), and CD68 (red).

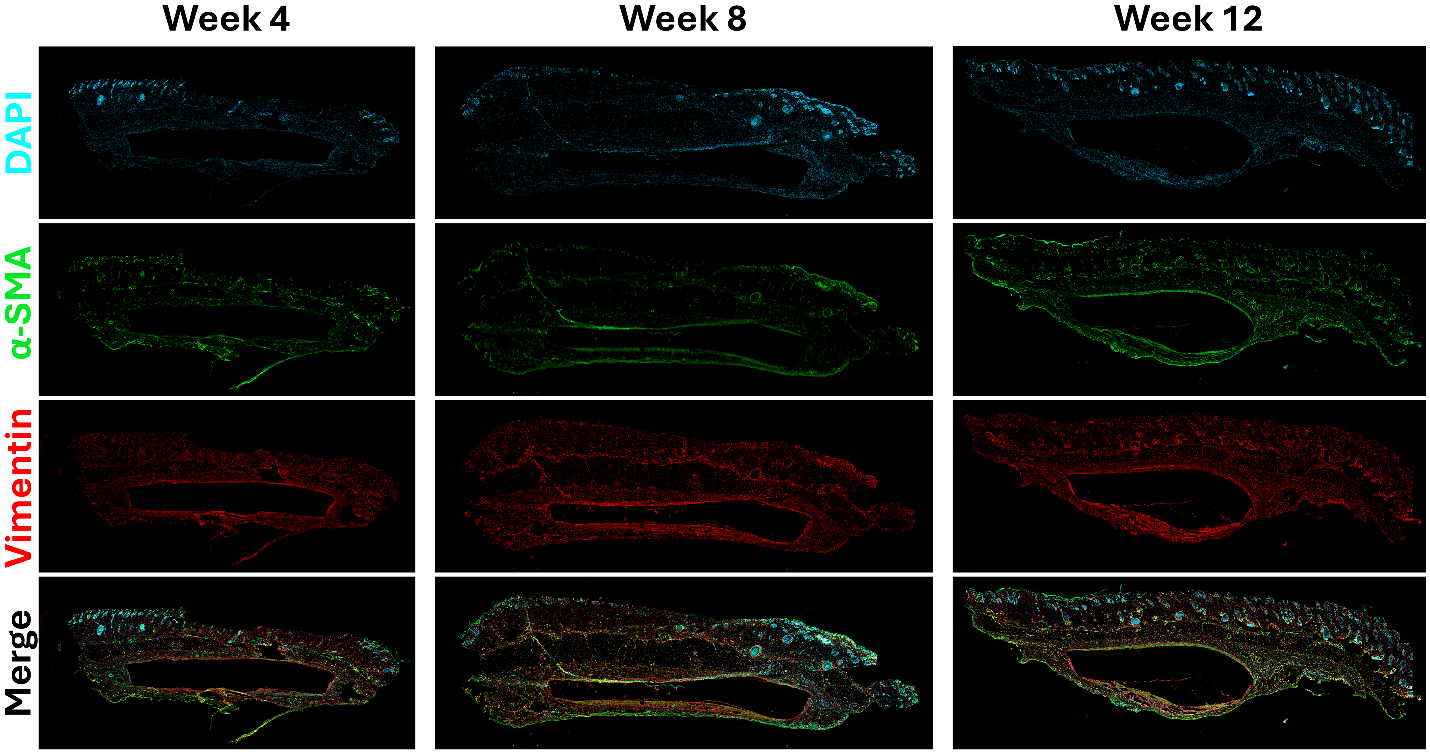

**Supplementary Figure S7.** Representative longitudinal sections from subcutaneous explants stained for DAPI (blue), α-SMA (green), and vimentin (red).

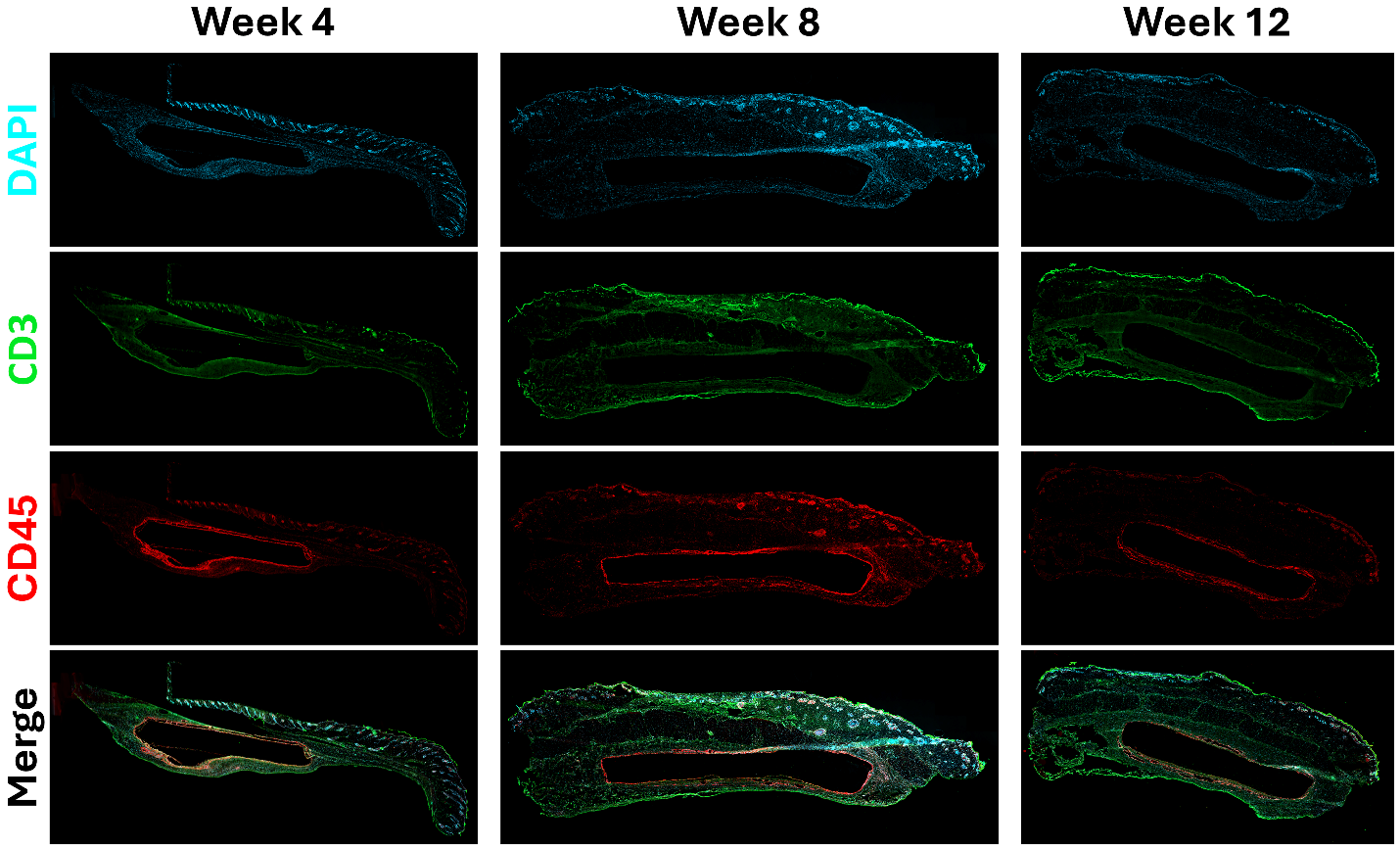

**Supplementary Figure S8.** Representative longitudinal sections from subcutaneous explants stained for DAPI (blue), CD3 (green), and CD45 (red).

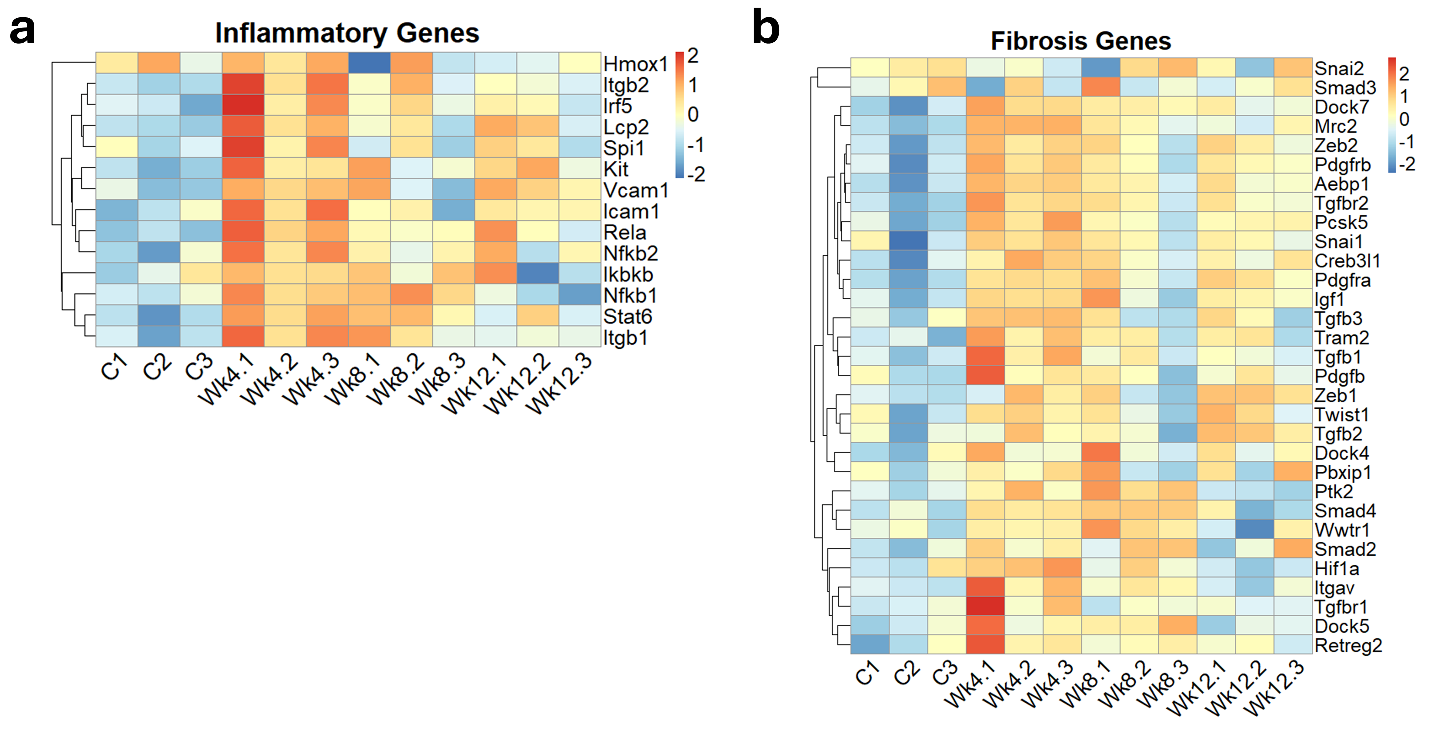

**Supplementary Figure S9.** Heatmaps displaying z-score normalized gene expression profiles for key markers associated with **(a)** inflammatory pathways and **(b)** ECM remodeling and fibrosis across all experimental groups.

**Video Captions:**

**Supplementary Video 1. Perfused DLP printed scaffold with internal channel.**

**Supplementary video 2. DLP printed tri-leaflet tissue engineered heart valve (TEHV) handling post printing.**

**Supplementary Video 3. Pulsatile flow testing of the DLP printed TEHV under pulmonary**

**condition.**

**Supplementary Video S4. Echochardiography of the DLP printed TEHV under pulmonary condition.**
